# Integrated transcriptomic analyses identifies host-targeting repurposing drugs for hepatitis C virus infection and related hepatocellular carcinoma

**DOI:** 10.1101/2025.05.17.654645

**Authors:** Sakshi Kamboj, Manoj Kumar

**Affiliations:** Virology Unit and Bioinformatics Centre, Institute of Microbial Technology, Council of Scientific and Industrial Research (CSIR), Sector 39A, Chandigarh-160036, India; Academy of Scientific and Innovative Research (AcSIR), Ghaziabad-201002, India

**Keywords:** Host-targeting, RNA-seq, drug repurposing, antiviral, anticancer

## Abstract

Hepatitis C virus (HCV) infection is a major risk factor in developing hepatocellular carcinoma (HCC). Drug development for HCV infection and HCC treatment is needed owing to treatment failure due to viral resistance and tumor heterogeneity. We analysed transcriptomic data of human liver tissues from multiple studies to identify differentially expressed genes in four study groups: (HCV+ vs Control); (HCV+HCC+ vs Control); (HCV+ AND HCV+HCC+ vs Control); and (HCV+HCC+ vs HCV+) respectively. We identified 597, 1321, 923, and 1526 upregulated and 474, 558, 336, and 989 downregulated genes for above respective studied groups. We performed functional, pathway enrichment, and protein-protein interaction analyses for identified upregulated genes. We identified and prioritized 39, 57, 58, and 110 repurposing drugs for 9, 17, 14, and 33 non-essential upregulated target genes based on drug group, action, and literature evidence. We also performed mutational analysis of key target genes. We found several dysregulated genes such as *CTLA4*, *GPC3*, *MDK*, *TUBB2B* etc. and many novel target genes like *CACNB4*, *HTR7*, *IGHG1*, *SLC22A12,* etc. We identified several potential repurposing drugs namely asenapine, cabergoline, dequalinium, epinastine, methysergide, loxapine, lurasidone, etc. This study identified prospective repurposing drugs using host-targeting approach for HCV infection and related HCC treatment.

## Introduction

Liver cancer is responsible for high global disease burden with ever-increasing incidence rate ^1^. About 0.8 million new cases of liver cancer are reported every year and are estimated to reach >1 million cases per year in the next decade ^2^. It is the fourth major cause of cancer-related deaths and sixth most prevalent cancer across the globe ^3^. Hepatocellular carcinoma (HCC) is the most common liver cancer accounting >90% cases ^4^. HCC is primarily developed as a result of chronic liver cirrhosis. Hepatitis C virus (HCV) infection is one of the major risk factors of liver cirrhosis. HCV is a pathogenic positive-sense single-stranded RNA virus which infects ∼58 million people across the globe with ∼1.5 million new infections every year ^5^ https://www.who.int/news-room/fact-sheets/detail/hepatitis-c. The incidence rate of HCC development is 1-4% in HCV infected individuals and 3-8% in chronic HCV patients ^6^.

Direct-acting antivirals (DAA) are used to treat HCV infection, achieving about 90% sustained virologic response. However, DAA therapy fails in about 8-10% of cases due to resistance-associated substitutions ^7^. So, due to resistance, treatment cost, availability, and persistence of HCC development; host-targeting agents (HTAs) are prospective options to the DAAs ^8^. Several studies have been conducted to explore the potential of HTAs in treating HCV infection. For instance, Hu et al., 2017 found an acyl coenzyme A: cholesterol acyltransferase inhibitor, avasimibe to be very effective in inhibiting the HCV infection ^9^. Bobardt et al., 2021 found that combination of cyclophilin and NS5A inhibitors inhibited the HCV infection *in vivo* without the development of resistance ^10^. Many HTAs have also been under clinical trials for activity against HCV like mirvirasen, alisporivir, erlotinib, bezafibrate. Thus, identification of potential HTAs for HCV treatment is promising and may revolutionize the field of HCV therapeutics. Likewise, For HCC management, different regimens based on the Barcelona Clinic Liver Cancer criterion are used and about 50-60% HCC patients are treated with systemic therapy ^4^. Currently used systemic regimens involve sorafenib, lenvatinib, cabozantinib, bevacizumab, regorafenib, tremelimumab etc.^11^. Since systemic therapy is administered depending on timing and tumour stage, there is a high failure rate, indicated by high mortality rate. Thus, there is always a need to look for new therapeutics for HCV-induced HCC.

As many key genes are reported to be modulated during HCV infection, along with the process of liver cirrhosis and ultimate HCC development. These key genes can be used as therapeutic targets for HCV infection as well as HCV-related HCC. For identifying these key modulated genes, analysing the differential gene expression during the diseased conditions can be helpful. Several computational studies have identified key target genes, miRNAs and pathways involved in different viral diseases, like Epstein-barr virus ^12^, Influenza virus ^13^. Moreover, many key genes can be prognostic markers for the development of HCC in HCV-infected patients. Some experimental and computational studies also reported differentially expressed genes (DEGs) during HCV infection and HCC ^14–16^. But there is a lack of integrative study which utilises the RNA-sequencing data from multiple studies to identify the DEGs during HCV infection and HCV-related HCC in clinical settings. In addition, no study has utilized these key genes as drug targets to identify repurposing drugs against HCV infection and HCV induced HCC.

In this study, we used the bulk RNA sequencing data of human liver tissues from HCV infected (HCV+) and/or HCV infected/induced HCC (HCV+HCC+) patients from multiple studies available in NCBI-sequence read archive (SRA). We used an in-house pipeline to identify the DEGs across different samples from different studies. We performed differential gene expression analysis for four study groups: (HCV+ vs Control), (HCV+HCC+ vs Control), (HCV+ AND HCV+HCC+ vs Control), and (HCV+HCC+ vs HCV+). As this study included samples from various studies, it is more robust in finding the common prevalent DEGs during HCV infection as well as HCV-induced HCC. These DEGs were subjected to functional and pathway enrichment analysis. The identified upregulated non-essential genes are used as targets to identify potential drugs for repurposing against HCV as well as HCV-induced HCC.

## Results

### Identification of differentially expressed genes during HCV infection and related HCC

We carried out differential gene expression analysis using the bulk RNA-sequencing data of human liver samples from four groups: (HCV+ vs Control), (HCV+HCC+ vs Control), (HCV+ AND HCV+HCC+ vs Control), and (HCV+HCC+ vs HCV+). The group HCV+ vs Control provided DEGs during the HCV infection as compared to the normal healthy condition. The group HCV+HCC+ vs Control identified the DEGs during HCV related HCC condition compared to normal healthy condition. While, HCV+ AND HCV+HCC+ vs Control group gives us DEGs during both HCV infection as well as HCV-related HCC condition. Likewise, the group HCV+HCC+ vs HCV+ identifies the DEGs specific for HCV-related HCC compared to HCV infection stage. The read counts for each study group are provided in **Supplementary Tables S2-S5.** Identified DEGs with p-adjusted value ≤ 0.01 and log2 fold change of ≥ 1.5 and -≤ 1.5 were taken as significant and selected for further analyses. For HCV+ vs Control, we identified 1072 significant DEGs with 597 upregulated and 474 downregulated genes. For HCV+HCC+ vs Control, we identified 1879 significant DEGs with 1321 upregulated and 558 downregulated genes. Similarly, we identified 1259 significant DEGs, with 923 upregulated and 336 downregulated genes for HCV+ AND HCV+HCC+ vs Control group. Likewise, for HCV+HCC+ vs HCV+ group, we identified 2515 significant DEGs with 1526 upregulated and 989 downregulated genes. The top five upregulated and downregulated genes for four study groups are given in **Table 1**. All significant DEGs for the four study groups are provided in **Supplementary Tables S6-S9**.

**Table 1.**
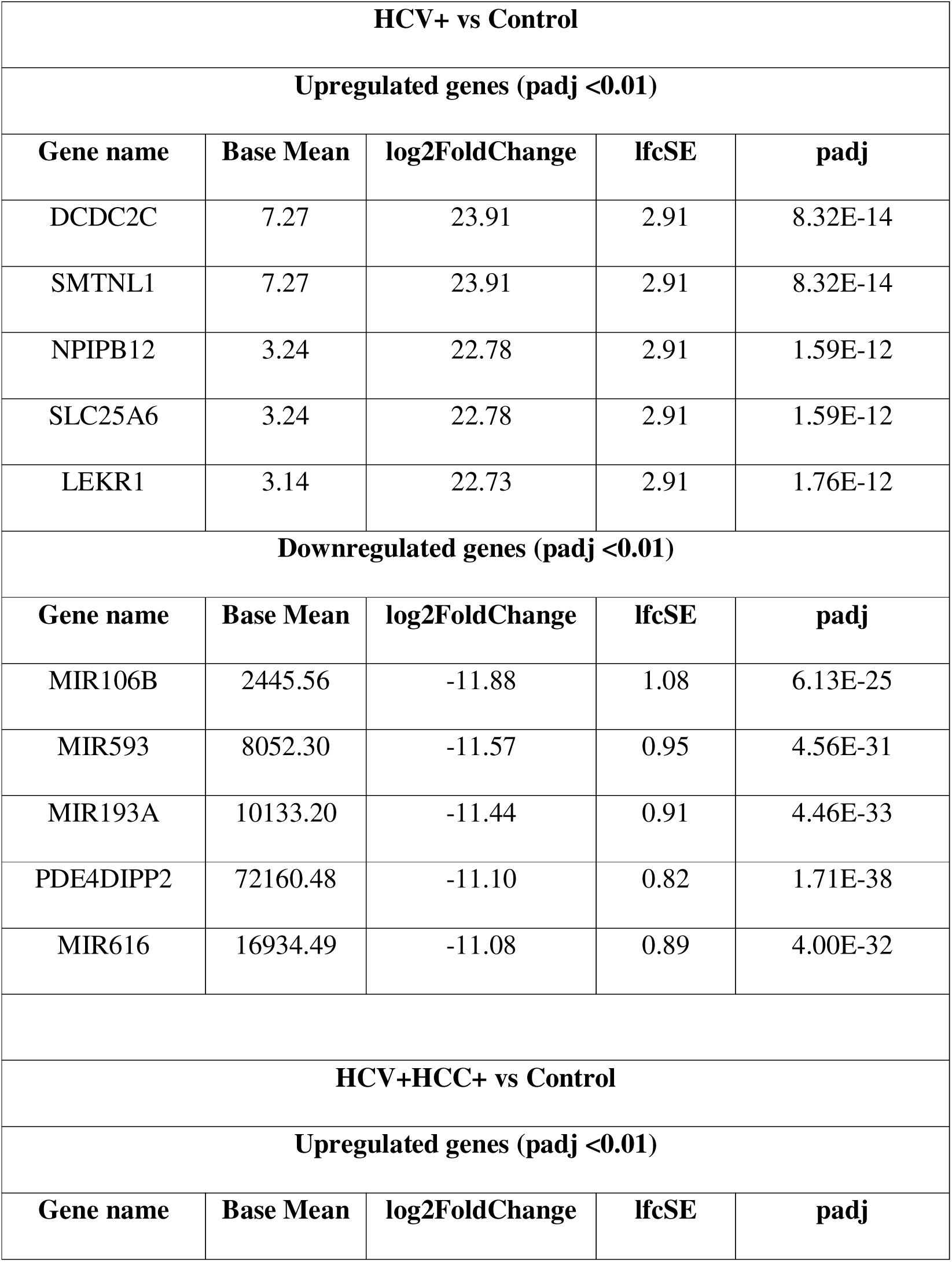
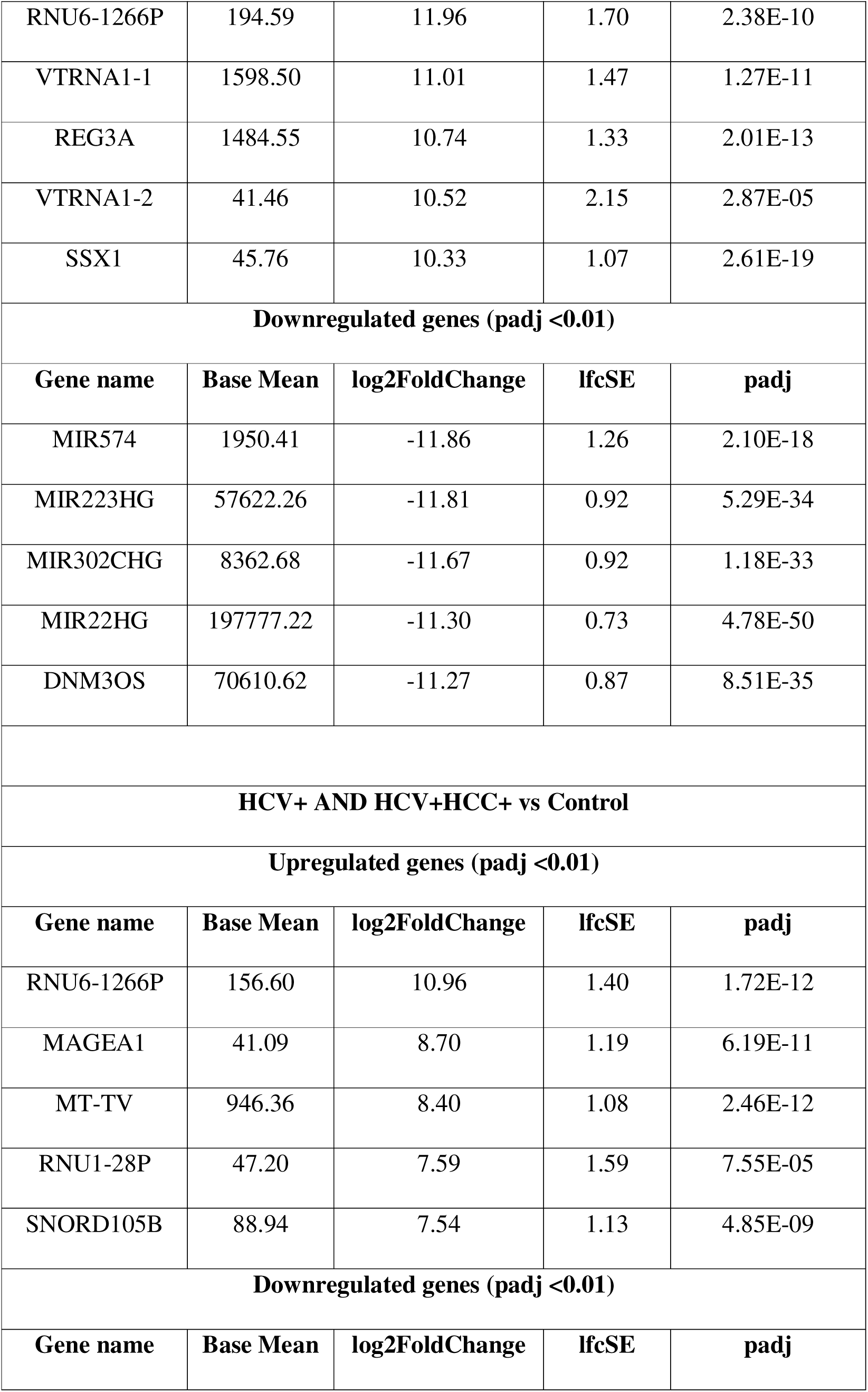
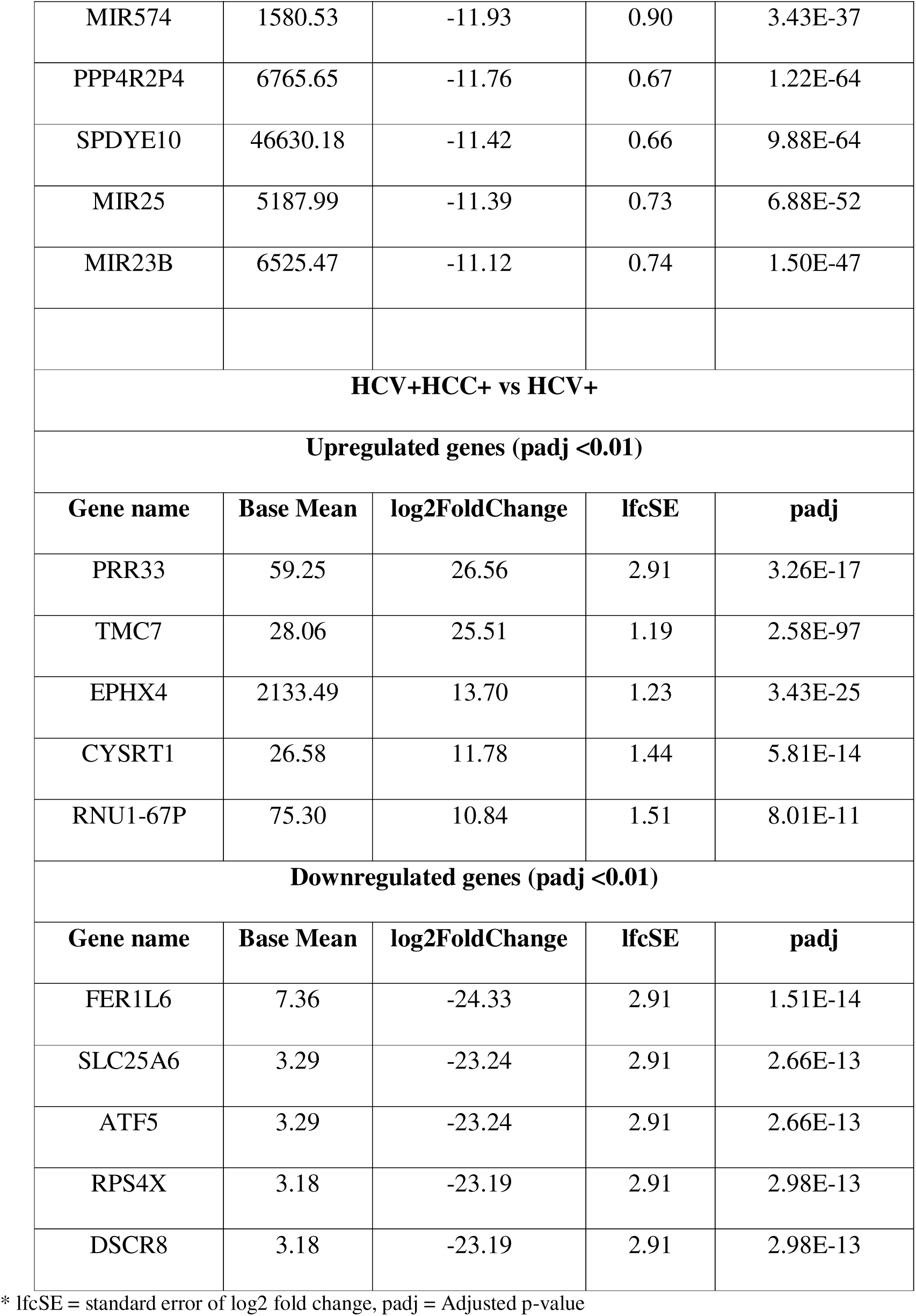
Table showing the significant top five upregulated and downregulated genes based on log2 fold change and adjusted p-value (padj) values in four study groups: (HCV+ vs Control), (HCV+HCC+ vs Control), (HCV+ AND HCV+HCC+ vs Control), and HCV+HCC+ vs HCV+:

The visualizations of DEGs for (HCV+ AND HCV+HCC+ vs Control) are shown in **Figure 1**. A scatter plot between log2FC and mean of normalised counts with p-adjusted value <0.05 called MA plot is shown in **Figure 1A**. This plot depicts the relationship of read counts with variability in expression of genes. The dispersion plot shows the dispersion of gene-wise estimates towards the fitted estimates, showing the final dispersion of data, given as **Figure 1B**. The overview of differentially expressed genes with log2FC and p-adjusted values are shown in the volcano plot in **Figure 1C**. The red dots denote the highly significant DEGs. For (HCV+ vs Control), (HCV+HCC+ vs Control), and (HCV+HCC+ vs HCV+) visualizations are given in **Supplementary Figure S1-S3** respectively.

**Figure 1.**
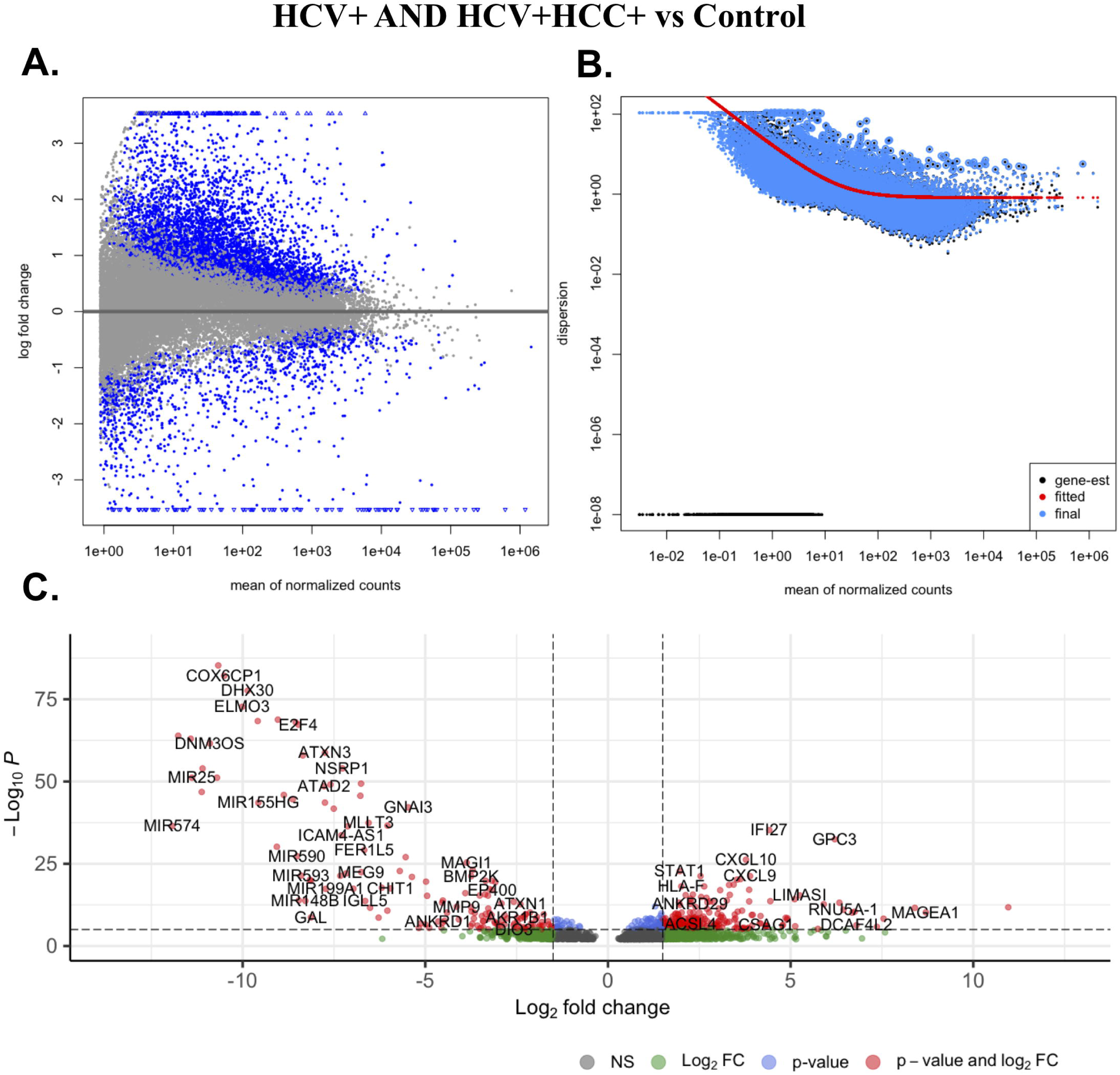
Visualizations of differential gene expression analysis of HCV+ AND HCV+HCC+ vs Control **A.** Mean-Average plot shows the relationship between the normalized counts and log2 fold change (log2FC) at p-adjusted value <0.05 **B.** Dispersion plot showing the gene wise dispersion estimates and fitted estimates **C.** Volcano plot showing the DEGs with padj < 0.01 and log2FC cut-off 1.5.

### Enriched gene ontology terms and pathways during HCV infection and related HCC

We performed Gene ontology (GO) and Kyoto Encyclopedia of Genes and Genomes (KEGG) enrichment analysis on the significant upregulated genes for all four studied groups. For HCV+ AND HCC+ vs Control group, highly enriched biological processes (BP) included the processes involved in immune response, immunoglobulin mediated immune response, RNA processing, response to virus, T cell receptor signalling pathway etc. The top 20 enriched BP for HCV+ AND HCC+ vs Control group are shown in **Figure 2A**. Likewise, most significant cellular components (CC) include immunoglobulin complex, nucleosome, extracellular region, external side of plasma membrane, etc. The top 20 enriched CC for HCV+ AND HCC+ vs Control are provided in **Figure 2B**. Enriched molecular functions (MF) belonged to categories: antigen binding, structural constituent of chromatin, protein heterodimerization activity, pre-mRNA 5’-splice site binding, etc. The top 20 enriched MF for HCV+ AND HCC+ vs Control are shown in **Figure 2C**. The enriched KEGG pathway belongs to systemic lupus erythematosus, neutrophil extracellular trap formation, alcoholism, viral carcinogenesis, hepatitis c, etc. The top ten enriched KEGG pathways for HCV+ AND HCC+ vs Control are shown in **Figure 2D**. Similarly, the top 20 enriched GO terms and KEGG pathways for HCV+ vs Control, HCC+ vs Control, and (HCV+HCC+ vs HCV+) are provided as **Supplementary Figure S4-S6** respectively. The detailed information on the enriched GO and KEGG pathways for four groups are provided in **Supplementary Tables S10 - S13**.

**Figure 2.**
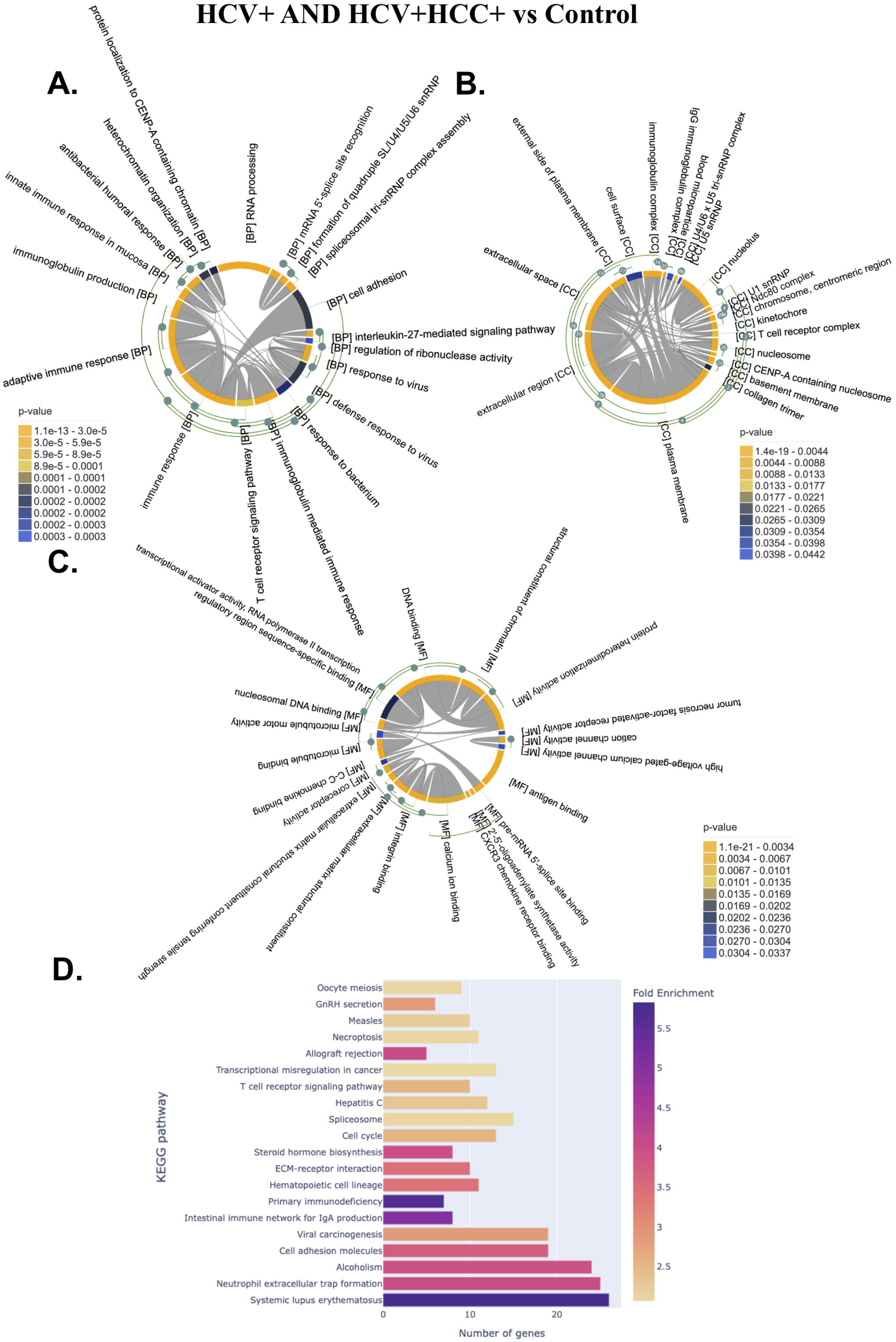
Enriched Gene Ontology terms and KEGG pathways for HCV+ AND HCC+ vs Control group upregulated genes. Top 20 Gene Ontology terms in **A.** Biological processes **B.** Cellular component **C.** Molecular Functions. **D.** Top 20 enriched KEGG pathways. For A-C, the green arcs on the outside parallel to the main chord diagram denote possible (hierarchical) clusters. The number on an arc-node represents the percentage of common genes between two nodes/clusters. The grey links inside the chord diagram connect pairs of GO term/clusters, and indicates the existence of common genes between them.

### Enriched hallmark gene sets during HCV infection and related HCC

Using hallmark gene sets from MSigDB, we identified enriched gene sets in diseased groups compared with control. For HCV+ AND HCV+HCC+ vs Control, hallmark gene sets such as interferon alpha response, interferon gamma response, G2M checkpoint, mitotic spindle, E2F targets, hedgehog signalling, etc. were enriched. For HCV+ vs Control, interferon alpha response, interferon gamma response, notch signaling, WNT/beta-catenin signalling, etc. gene sets were enriched. Likewise, for HCV+HCC+ vs Control, G2M checkpoint, E2F targets, mitotic spindle, interferon alpha response, etc. Similarly, For HCV+HCC+ vs HCV+, mitotic spindle, G2M checkpoint, E2F targets, unfolded protein response, MYC targets, etc. are enriched. The GSEA output for each study group is provided in **Supplementary Tables S14**. We also checked the distribution of upregulated genes into different gene families available in MSigDB. For HCV+ AND HCV+HCC+ vs Control, we found that out of 923 upregulated genes, 92 belong to major gene families. These genes were categorized into eight different gene families i.e., cytokines and growth factors (21), transcription factors (11), homeodomain proteins (16), cell differentiation markers (31), protein kinases (9), translocated cancer genes (13), oncogenes (13), and tumor suppressors (2). The distribution of (HCV+ AND HCV+HCC+ vs Control) upregulated genes to different gene families are shown in **Figure 3**. Likewise, for HCV+ vs Control 597 upregulated genes, 55 genes belong to eight major gene families: cytokines and growth factors (9), transcription factors (7), homeodomain proteins (4), cell differentiation markers (29), protein kinases (4), translocated cancer genes (5), oncogenes (6), and tumor suppressors (1). Similarly, for HCV+HCC+ vs Control 1321 upregulated genes, 174 genes belong to eight different gene families as follows: cytokines and growth factors (34), transcription factors (74), homeodomain proteins (26), cell differentiation markers (21), protein kinases (31), translocated cancer genes (17), oncogenes (17), and tumor suppressors (11). For HCV+HCC+ vs HCV+, 1526 upregulated genes, 131 genes belong to eight different gene families as follows: cytokines and growth factors (28), transcription factors (25), homeodomain proteins (31), cell differentiation markers (11), protein kinases (28), translocated cancer genes (14), oncogenes (14), and tumor suppressors (7). The distribution of upregulated genes from HCV+ vs Control, HCC+ vs Control, and (HCV+HCC+ vs HCV+) into different gene families are shown in **Supplementary Figure S7-S9** respectively. The genes belonging to different gene families for each group are provided in **Supplementary Table S15**.

**Figure 3.**
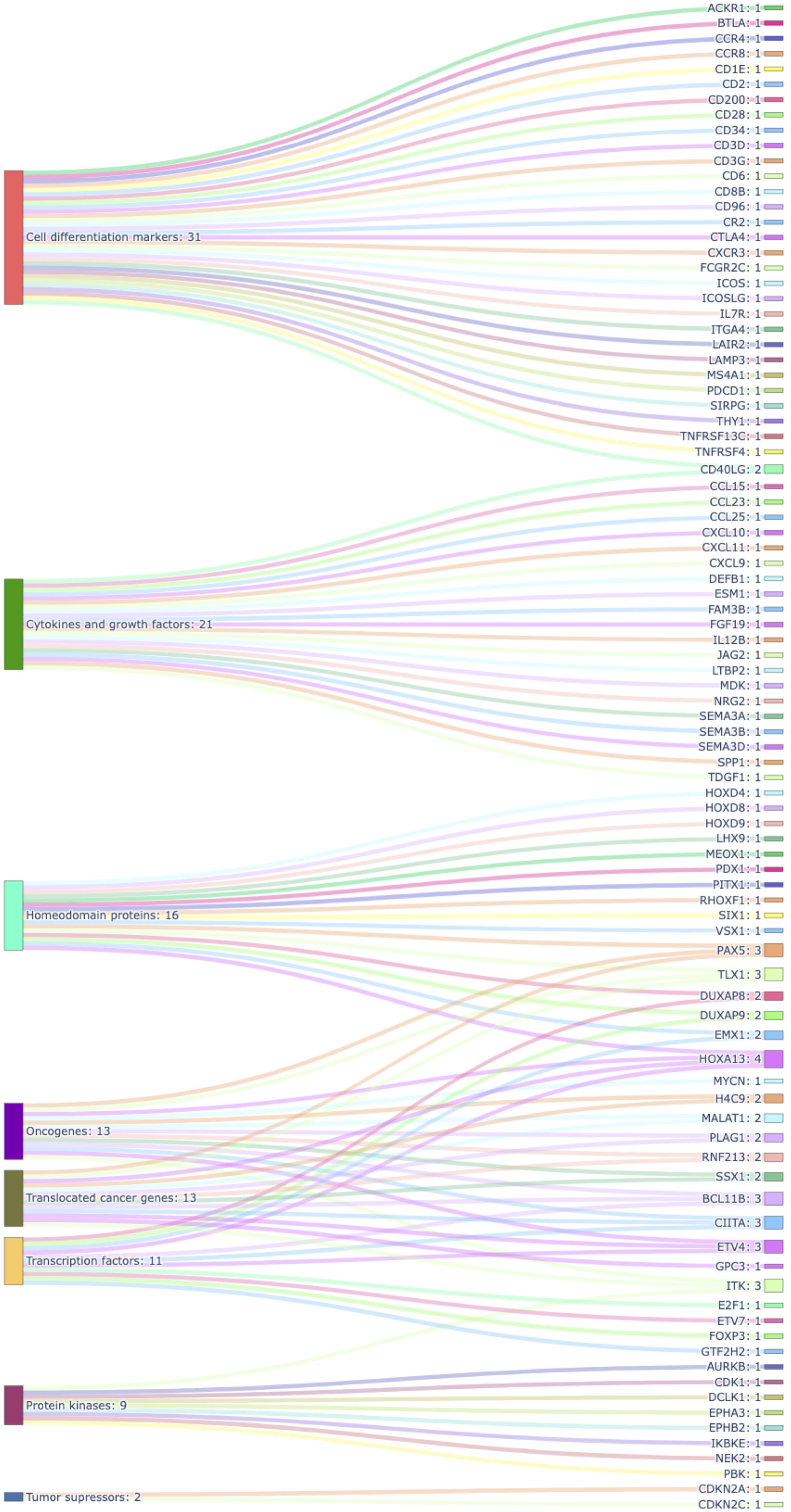
Distribution of upregulated genes from HCV+ AND HCV+HCC+ vs Control group into different gene families in Sankey plot. The number written next to each gene family indicates the number of genes belonging to that gene family. The number next to gene names denotes the number of gene families to which that gene belongs.

### Protein-protein interaction networks revealed the interacting proteins with upregulated genes

The significantly upregulated genes from four groups were analysed for their protein-protein interaction (PPI). We used the STRING database plugin in Cytoscape v3.9.1 software for PPI analysis. PPI network for HCV+ AND HCV+HCC+ comprises 969 nodes and 8811 edges, shown in **Figure 4**. For HCV+ vs Control, PPI network contains 307 nodes and 1129 edges, shown in **Supplementary Figure S10**. PPI network for HCC+ vs Control comprises 577 nodes and 2865 edges, depicted in **Supplementary Figure S11**. PPI network for HCV+HCC+ vs HCV+ comprises 1064 nodes and 7664 edges, given in **Supplementary Figure S12**.

**Figure 4.**
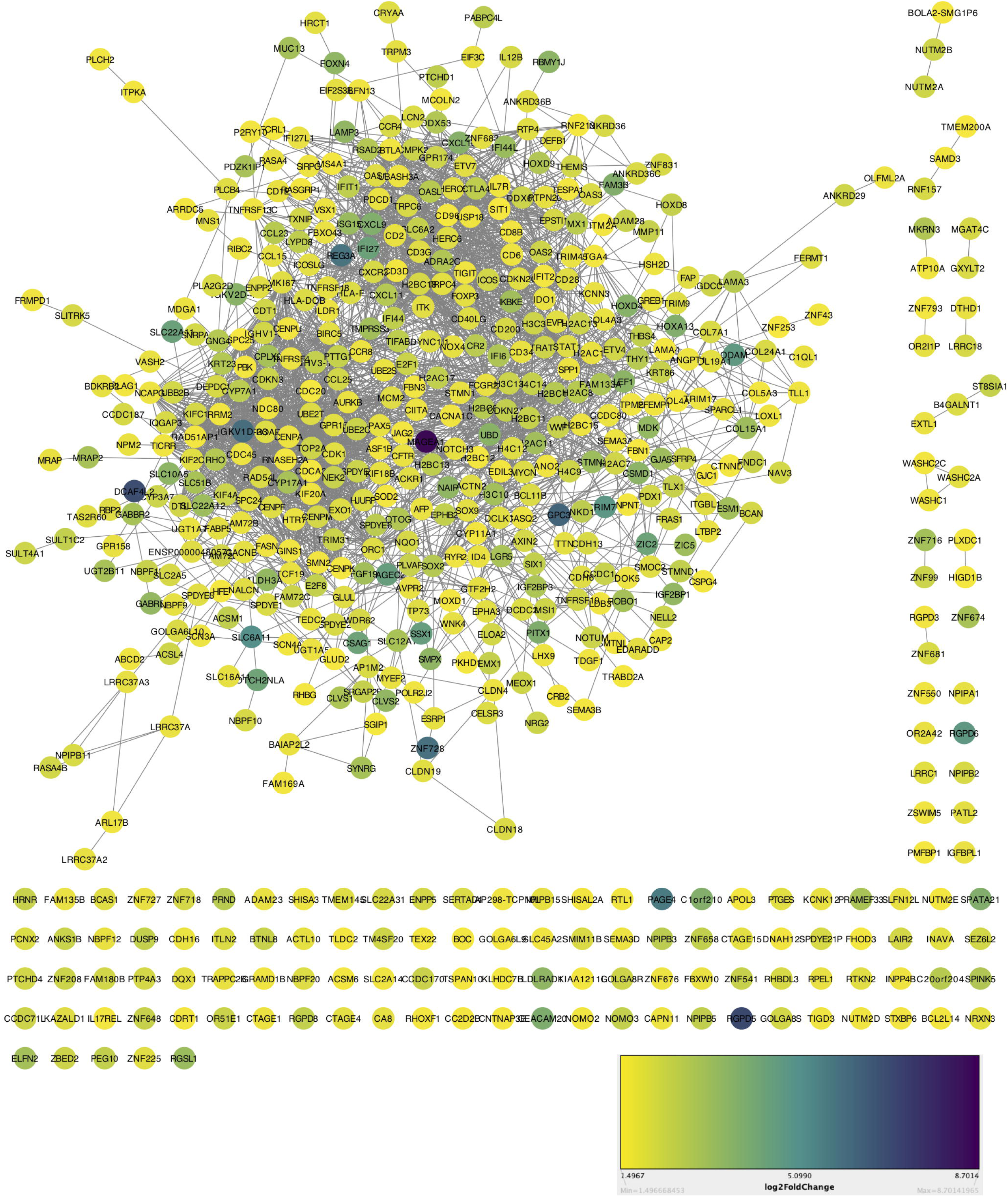
Network showing protein-protein interactions of significant upregulated genes in HCV+ AND HCV+HCC+ vs Control. The node colors are shown on the basis of log2 fold change provided in legend.

### Essential and non-essential genes from upregulated genes

We analysed the 923 upregulated genes from HCV+ AND HCV+HCC+ vs Control and identified 682 non-essential genes. For HCV+ vs Control, HCV+HCC+ vs Control, and HCV+HCC+ vs HCV+, out of 597, 1321 and 1526 upregulated genes, 500, 886, and 1049 genes belong to non-essential genes category respectively. The non-essential genes for each group are provided in **Supplementary Table S16**.

### Identification of repurposing drugs targeting upregulated non-essential genes

We used the upregulated non-essential genes from four groups as drug targets for identifying drugs. We looked into the ‘DrugBank’ repository for the drugs against these gene targets. We found 215, 486, 682, and 924 drugs against 31, 76, 44, and 109 target genes for (HCV+ vs Control), (HCV+HCC+ vs Control), (HCV+ AND HCV+HCC+ vs Control), and (HCV+HCC+ vs HCV+) groups respectively. We prioritised and selected the potential repurposing drugs, based on the information such as not previously tested for these disease conditions, drug groups (approved, approved; investigational, approved; experimental, approved; vet approved), drug action (antagonist, inhibitor, antibody, blocker, modulator, etc.). We selected 39, 57, 58, and 110 drugs for 9, 17, 14, and 33 target genes for the above groups respectively as potential repurposing drugs candidates. The steps followed to identify, select and prioritise the target genes and potential repurposing drug candidates along with their number for the four study groups is given as **Figure 5**. The number of overlapping target genes for all groups are shown as Venn diagram in **Supplementary Figure S13**. The list of all identified repurposing drugs with other information for four groups are given in **Supplementary Tables S17 - S20**. The prioritised repurposing drugs for four groups are shown in a Sankey plot with different categories in **Figure 6**.

**Figure 5.**
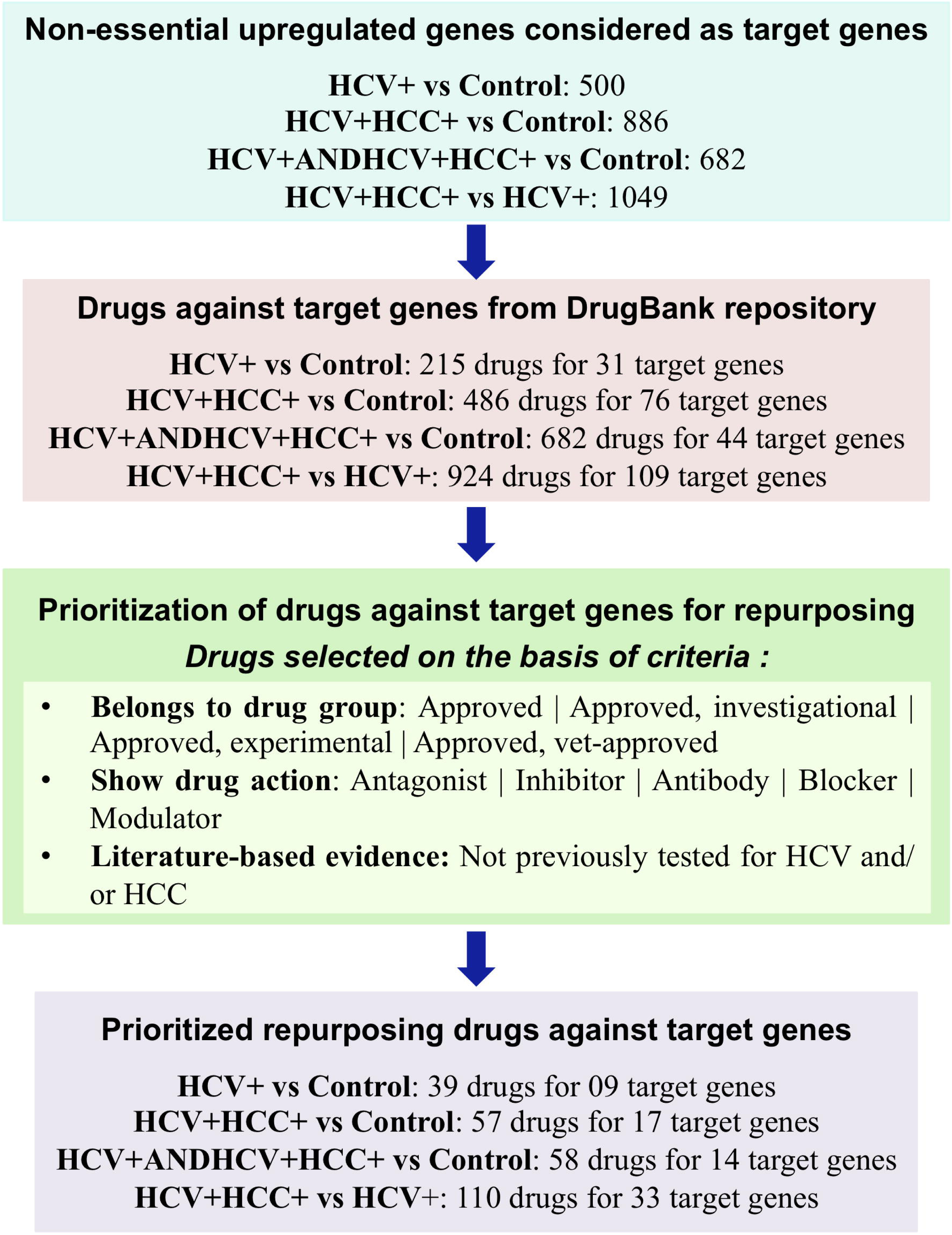
Flow diagram showing the steps involved in the identification of drug target genes and their drugs from the DrugBank repository for the four study groups. The prioritization is given to the drugs meeting all three criteria (in bullet points) and are selected as prioritized repurposing drug candidates along with their target genes.

**Figure 6.**
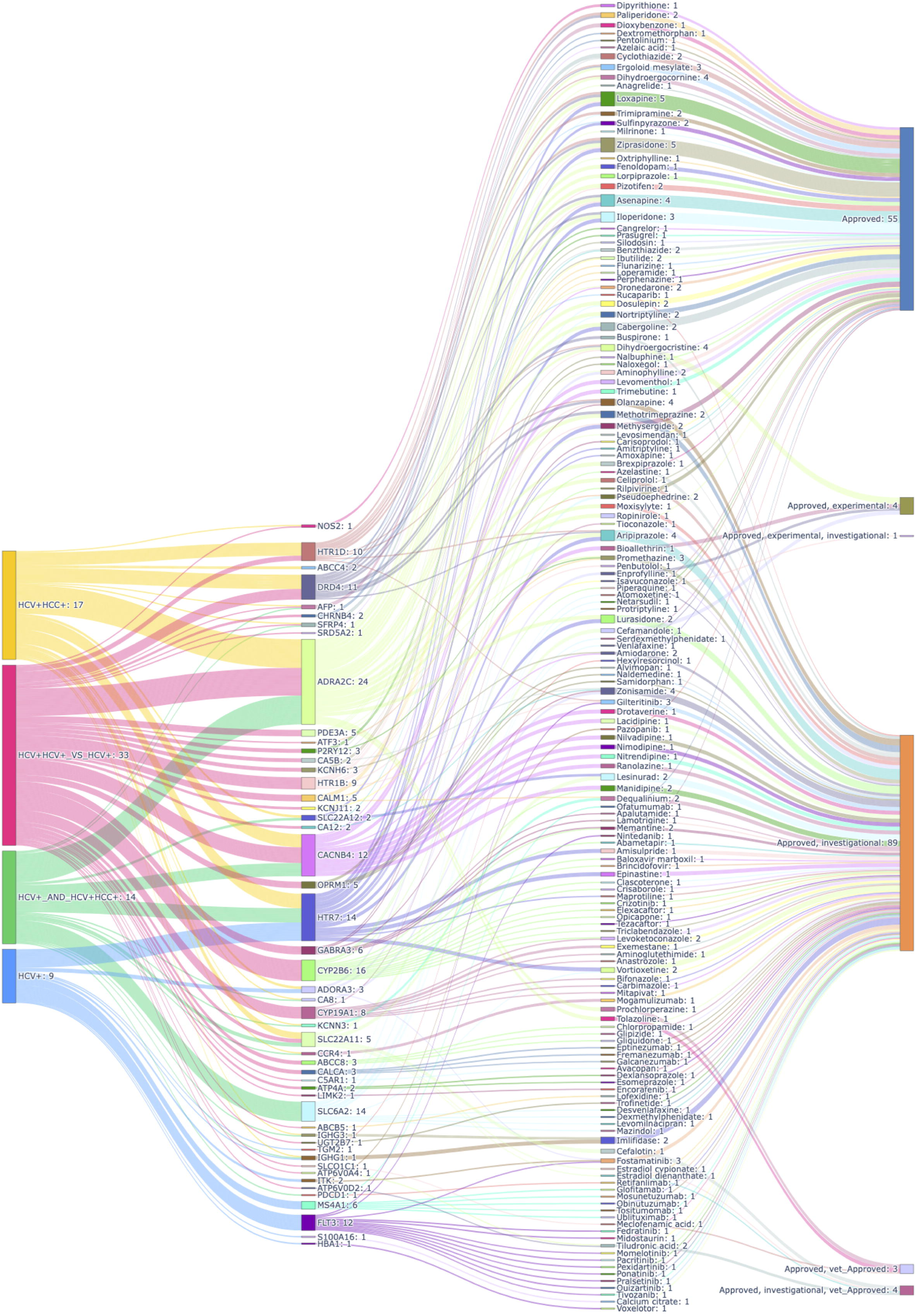
Prioritised repurposing drugs with their target genes and drug groups for study groups: (HCV+ vs Control), (HCV+HCC+ vs Control), (HCV+ AND HCV+HCC+ vs Control), and (HCV+HCC+ vs HCV+). The numbers written with the study groups (leftmost panel) indicate the number of prioritised target genes identified for respective study group. For the numbers next to target genes are the number of prioritised drugs identified for each target gene (second to left). For numbers next to drug names are the number of target genes for each drug (second to right). The numbers next to drug groups are the number of drugs belonging to each drug group (Rightmost panel).

### Mutational profile of prioritised target genes for drug repurposing

As mutations in the target genes can lead the development of resistance towards the drugs against particular target genes, we checked the mutational profile of the identified target genes for each study group. The mutational profiles of the target host genes of identified potential repurposing drugs for study groups were assessed using the mutational data of eight different studies available in cBioPortal (https://www.cbioportal.org/). These eight studies were mainly conducted for hepatocellular carcinoma patients and included 2589 samples and 2538 patients in total. We identified mutations, copy number alterations and structural variants for the target genes. The mutational profile for unique prioritised target genes (50 in number, median mutation rate: 0.8%) from four study groups is shown in **Figure 7**. The mutational profile of target genes for four study groups are shown in **Supplementary Figures S14-S17** respectively. The target genes showed very few mutations with a median of 0.6% for HCV+ vs Control, 0.7% in HCV+HCC+ vs Control, 0.6% in HCV+ AND HCV+HCC+ vs Control, and 0.9% in HCV+HCC+ vs HCV+ group. The percentage mutation rate of target genes with drugs are given in **Supplementary Table S21**. Result of mutational analysis further indicates that these prioritised genes are good targets for drug repurposing.

**Figure 7.**
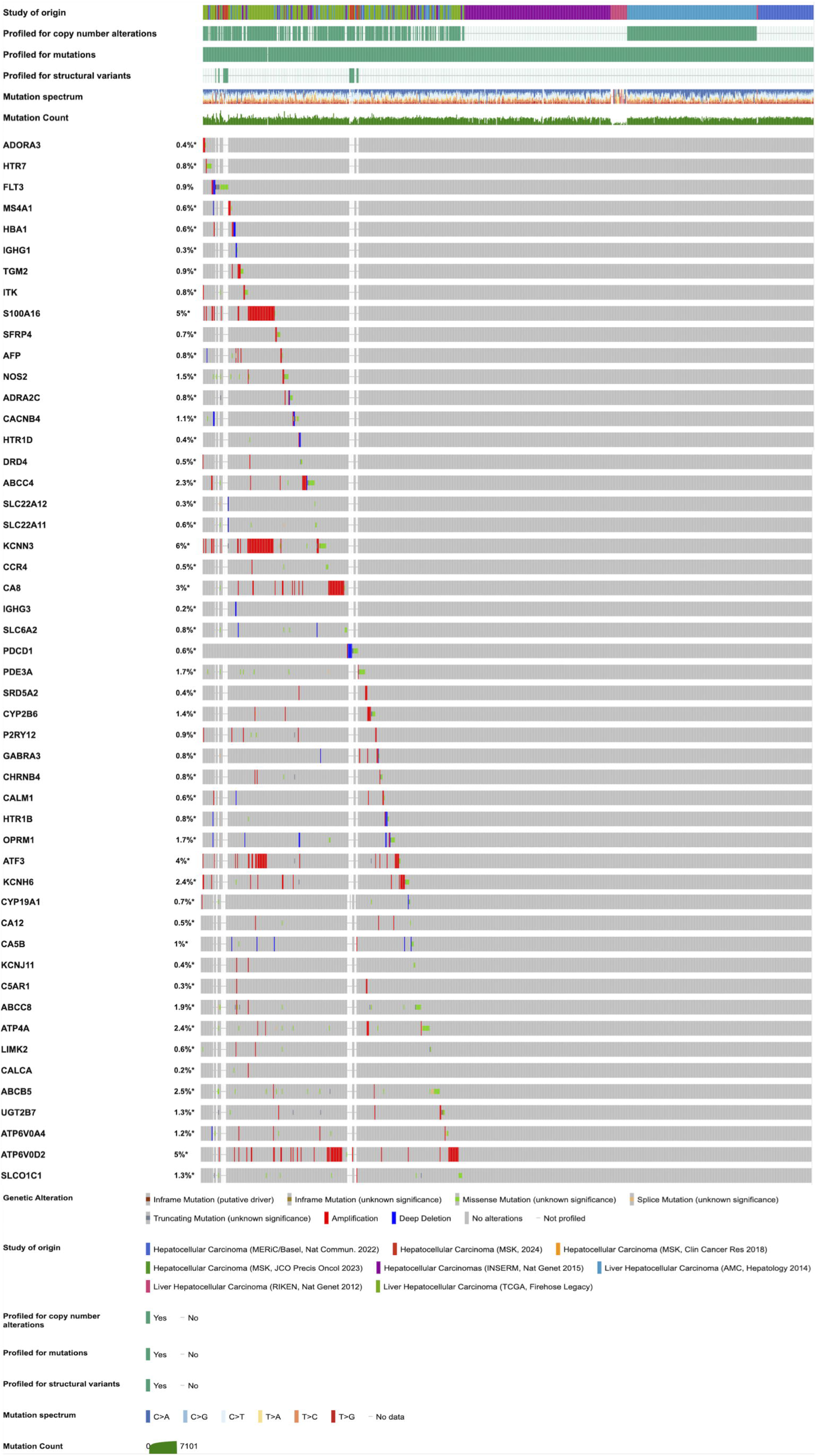
Oncoprint image showing the mutational profiles of prioritised unique target genes for all four groups: (HCV+ vs Control), (HCV+HCC+ vs Control), (HCV+ AND HCV+HCC+ vs Control), and (HCV+HCC+ vs HCV+).

### Predicted siRNAs and sgRNAs for prioritised target genes

We also explored the nucleic acid intervention tools namely siRNAs and sgRNAs as therapeutic agents against the prioritized target genes. We identified the siRNAs and sgRNAs for the unique prioritized target genes (50 in number). We predicted siRNAs for prioritized target genes using siRNApred web server The 10 predicted siRNAs with their efficacies for 48 target genes are given in **Supplementary Table S22**. We also carried out designing of sgRNAs to employ the CRISPRi mechanism of silencing the transcription of targeted genes. We designed sgRNAs for 50 prioritized target genes with maximum on-target and minimum off-target activity. The 10 designed sgRNAs for enzyme SpyoCas9 for each target gene are provided in **Supplementary Table S23**.

## Discussion

In this study, we used RNA-sequencing data of human liver tissues from multiple studies from patients with HCV infection or HCV related HCC condition in clinical settings along with healthy liver samples. This data was used to carry out integrative and combinatorial analysis of differential gene expression for four study groups: HCV+ vs Control, HCV+HCC+ vs Control, HCV+ AND HCV+HCC+ vs Control, and HCV+HCC+ vs HCV+. These comparisons provided DEGs during HCV infection, HCV related HCC, HCV infection with HCV-related HCC from healthy control and HCV-related HCC compared to HCV infection.

In study group HCV+ vs Control, we identified 1072 significant DEGs comprising 597 upregulated and 474 downregulated genes during HCV infection. Among these upregulated genes, 500 genes were found to be non-essential genes. Many DEGs from our study are also reported in previous studies. For instance, genes like *STAT1*, *OAS1*, *OAS2*, are reported to be dysregulated during early stages of liver fibrosis in chronic HCV patients ^17^. Similarly, higher expression of genes *NRIR, BISPR, RSAD2* are reported to be associated with chronic HCV infection ^18^. In addition to reported genes, we found several novel genes dysregulated during HCV infection. Genes like *ZBED1*, *SLC25A6*, *ZNF683*, *MIR199A1*, *S100A12*, *CALCA* etc. showed highly up or down regulated expression **(Supplementary Table S6)**. Some of the novel genes were also found to be associated with other infections or viral diseases. For example, *IGHV1-18*, *IGHV3-13*, *IGLV3-23, IGLV3-21,* ^19^ were found to be dysregulated during severe acute respiratory syndrome coronavirus 2 (SARS-CoV-2) infection and tested for effectiveness of blockage in therapeutic intervention. Other novel genes reported were known to be dysregulated in other viral infections like *ADAM28* for Human papilloma virus ^20^, *PLA2G2D* for Epstein-barr virus ^21^, etc. We identified 215 drugs for 31 genes from 500 non-essential genes from “DrugBank ‘’ repository. Some identified drugs are also reported for their activity against HCV infection, showing the robustness and reliability of our study. *TLR7* agonists, isatoribine ^22^ and resiquimod ^23^ were subjected to phase 1 and two double-blind, placebo-controlled phase 2a trials respectively for the treatment of HCV infection. *IDO1* inhibitory drug, cannabidiol was found to exhibit the inhibitory activity against HCV ^24^.

Likewise for the HCV+HCC+ vs Control group, we found 1879 significant DEGs with 1321 upregulated and 558 downregulated genes. Among these upregulated DEGs, 886 non-essential genes were identified. Several genes found in our study were reported for HCC in literature. Genes such as *CCNB1*, *CCNE1*, *E2F1*, *EZH2*, *MCM4*, *PBK*, and *RRM2* were reported to be highly expressed during HCC progression, making them to be potential HCC biomarkers other than AFP ^25^. We found high expression of human vault RNA, *VTRNA1-1* in our study that coincided with study by Ferro et al., 2022 in which they explored the role of *VTRNA1-1* in HCC and associated resistance to apoptosis ^26^. We also reported several novel genes highly dysregulated during HCV related HCC like *RNU6-1266P*, *VTRNA1-2*, *RNU5B-1*, *RNU1-11P*, *SNORD71*, *MIR574*, *MIR199A1*, *ADGRA1*, etc. **(Supplementary Table S7).** From the identified novel genes, we found some genes are reported for other carcinomas also. Wu et al., 2022 showed that targeting *HTR1D* inhibits the proliferation and progression of pancreatic cancer ^27^. Similarly, *HSD3B2* for adrenocortical carcinoma ^28^, and prostate cancer ^29^ is reported. We also identified 486 drugs for 76 target genes from 1321 non-essential genes. Among these drugs, some are also reported to be administered for HCC management. For instance, trilostane, an *HSD3B2* inhibitor, was found to inhibit HCC growth in a dose-dependent manner ^30^. Zolmitriptan, an *HTR1D* agonist, was reported to activate caspase mediated apoptosis, inhibiting HCC growth ^31^.

For HCV+ AND HCV+HCC+ group, we identified 1259 significant DEGs consisting of 923 upregulated and 336 downregulated genes. We found 682 genes belong to non-essential genes category from the upregulated genes. Several genes and their expressions from our study also match with the findings from other studies. For example, Genes like *CCR4* ^32^, *HRNR* ^33^ etc. are reported to be dysregulated during HCV infection as well as HCC condition. Gene *GPC3* is found to be associated with HCV-related HCC progression and can also be used as biomarker for HCV related HCC ^34^. Similarly, IFN-stimulated genes such as *IFI27*, *IFIT1*, *IFI6*, *ISG15*, and *CXCL10* shows gene overexpression during HCV infection ^35^ along with *IFI27* for HCC ^36^. Along with previously reported genes, we also found certain novel genes including prioritised genes which are dysregulated during HCV infection to HCC progression. Genes such as *RNU6-1266P*, *RNU5B-1, ZNF728*, *MIR574*, *MIR199A1*, *SPDYE10*, *CEACAM5* etc. are highly dysregulated novel genes from HCV infection to HCC proliferation **(Supplementary Table S8)**. Few of these novel genes are also reported to be involved in other viral infections and cancers. For instance, Wong et al., 2022 reported *PLA2G2D* involved in eicosanoid signalling as a good therapeutic target for SARS-CoV-2 infection ^37^. Kim et al., 2022 found that overexpression of calsequestrin 2 (*CASQ2*) increases tumorigenesis in breast cancer ^38^. Serotonin receptors including *HTR7* reported to be overexpressed during gastric cancer ^39^. We also identified 682 drugs for 44 target genes from 682 non-essential genes from “DrugBank’’ repository. Some identified drugs are also reported for antiviral and anticancer activity against HCV infection and HCC condition. For instance, in a pilot clinical trial, tremelimumab, *CTLA4* inhibitor was reported to have antiviral and antitumor activity for chronic HCV and HCC conditions ^40^. Tislelizumab, *PDCD1* antibody, reported to show antitumor activity for HCC and tested in a phase 2, open, non-randomized trial for advanced HCC ^41^. Sertraline, an *SLC6A2* inhibitor, was tested in combination with simvastatin for antiviral activity against HCV in phase 1b pilot study. It was also studied in combination with sorafenib for anti-HCC activity ^42^.

For the HCV+HCC+ vs HCV+ group, we identified 2515 significant DEGs comprising 1526 upregulated and 989 downregulated genes. Among 1526 upregulated genes, 1049 were non-essential genes. This study group provided the genes involved in the development and progression of HCC from HCV infection. Many genes from our study are reported to be associated with HCC tumorigenesis in previous studies. For instance, gene *GLMP* is reported to be involved in lipid metabolism in NAFLD-associated HCC progression ^43^. Similarly, gene *MAGEC2,* with higher expression in HCC tissues, is implied as a novel prognostic marker and therapeutic target for HCC ^44^. Likewise, gene *FANCG* is found to be upregulated in non-alcoholic steatohepatitis associated HCC than alcoholic hepatitis related HCC ^45^. Genes like *ZIC2* ^46^, *TINAG* ^47^, etc. promote HCC progression. Other than previously reported genes, we also identified several novel genes which are dysregulated during HCC condition progression from HCV infection. Some such genes are *PRR33*, *CYSRT1*, *RNU1-67P*, *LINC01058*, *SLC25A6*, *RPL27AP5*, *SNX19P4*, *RPL21P28* etc **(Supplementary Table S9).** Some of these novel genes are also reported for other cancers like *PAGE4, KBTBD11* in colorectal cancer ^48, 49^, *SNORA66* in diffuse large B-cell lymphoma ^50^. In addition, we identified 924 drugs for 109 target genes from 1049 non-essential upregulated genes. Some identified drugs are also reported previously against HCC progression. *UGT2B7* substrate, dabigatran etexilate along with its derivatives were found to be effective against HCC tumor cells proliferation by inhibiting thrombin activity ^51^. *OPRM1* antagonist, butorphanol is reported to inhibit metastasis and angiogenesis in HCC by regulating *MAPK* signalling ^52^.

Functional and pathway enrichment analysis for each study group showed the enriched pathways and associated functions during the diseased conditions. We also found that the prioritised target genes belonged to certain enriched GO terms and KEGG pathways. Several identified enriched pathways are also previously reported to be involved in HCV infection. For instance, GO terms: immune response, extracellular region are reported during HCV infection ^53, 54^. Additionally, immune responses are also targeted for the HCV-induced fibrosis treatment ^55^. While, involvement of KEGG pathways: T cell receptor signalling pathway, PD-L1 expression and PD-1 checkpoint pathway during HCV infection are also reported ^56^. For HCV+HCC+ vs Control group, Many identified enriched GO terms and KEGG pathways are also previously reported for HCC. For example, enriched extracellular matrix related GO terms (GO:0030198, GO:0005576, GO:0005201) coincides with involvement of extracellular vesicles in HCC development ^57^. Extracellular vesicles are also studied extensively for HCC treatment and diagnosis ^58^. KEGG pathway: viral carcinogenesis can be directly related with the involvement of HCV in development of HCC.

Similarly HCV+ AND HCV+HCC+ vs Control group showed common enriched GO terms and KEGG pathways between HCV infected tissue and HCV related HCC tissue. For instance, T cell receptor signalling pathway is reported to be involved during the HCV infection ^56^ as well as in HCC condition ^59^. Similarly, Bergqvist et al., 2003 reported the modulation in T cell response upon HCV infection by T cell receptor induced Ca2+ oscillations which can be related with our finding of enriched GO term calcium ion binding ^60^. Lima et al., 2018 found that the cell adhesion molecules are co-expressed with the known biomarker *AFP* during HCC, linking it with enriched KEGG pathway: Cell adhesion molecules ^61^. Likewise, for HCV+HCC+ vs HCV+, Several enriched GO terms and KEGG pathways are found to dysregulated during the HCC condition irrespective of HCV infection. Sanchez et al., 2021 showed alterations in retrograde endocannabinoid signalling pathway from the lipidomic profiling of plasma exosomes in HCC patients with cirrhosis ^62^. Zhou et al., 2022 reported the enrichment of cell cycle genes from The Cancer Genome Atlas data and role of SNHG1-centered ceRNA network in cell cycle regulation in HCC ^63^.

We found that some enriched pathways also differed from previous findings but these findings can be related to extrahepatic manifestations due to HCV infection and HCV-related HCC. For instance, HCV is suspected to be the trigger for connective tissue disease, systemic lupus erythematosus^64^. Vucur et al., 2023 reported that necroptosis signalling can promote liver cancer development ^65^. Our findings also coincide with some proteomics studies like many genes coding for proteins dysregulated in our study are also identified previously. Genes *ISG15*, *IFIT1* upregulated in our study are also reported to be overexpressed during HCV infection in previous proteomic study by Broering et al., 2016 ^66^. Similarly, genes *A2MP1*, *APOA4* ^67^, *CXCL9* ^68^ etc. during HCV infection, while *IGLC2*, *IGKC* in HCV-related HCC ^69^, etc. are also dysregulated in our study. We also predicted and designed siRNAs and sgRNAs against the prioritized target genes. In addition, lower is the mutation rate of the target genes, lower will be the chances of development of resistance towards the drugs against the target genes. Mutational analysis of prioritised target genes showed low mutation rate, evident to potential of identified genes as good targets for drug repurposing.

We have carried out this study using the raw transcriptomics data from multiple studies analysed by in-house pipeline. However, there are some limitations of our study. Even though there is large availability of RNA-sequencing data from non-clinical/cell culture and clinical human blood (PBMCs) samples from HCV infection studies. We preferred for the human liver samples in the clinical settings for both HCV infection and related HCC. We also only select studies having the required metadata like infection status, drug therapy, tumour/non-tumour status, etc for our analysis. We could not integrate proteomics data with transcriptomics data due to non-availability of desired proteomic data from human liver samples. As this is purely bioinformatics study, the findings can be further experimentally validated in the future studies. In conclusion, this integrative and combinatorial study helped in finding the key genes dysregulated both during HCV infection to HCC progression. These key genes are used as targets to identify the potential repurposing drugs to be used as HTAs for treatment of HCV infection and HCC condition. We identified several novel dysregulated genes as well as potential repurposing drugs for HCV infection and HCC progression. This study assists in finding new drugs for HCV infection in view of emerging drug resistance to existing DAAs. Furthermore, many of the proposed repurposing drugs identified in our analyses are shown to have experimental evidence at clinical/preclinical stages for HCC management.

## Methodology

The overall workflow of this study is shown in **Figure 8**. The detailed methodology followed for conducting this study is given in **Supplementary File 1**.

**Figure 8.**
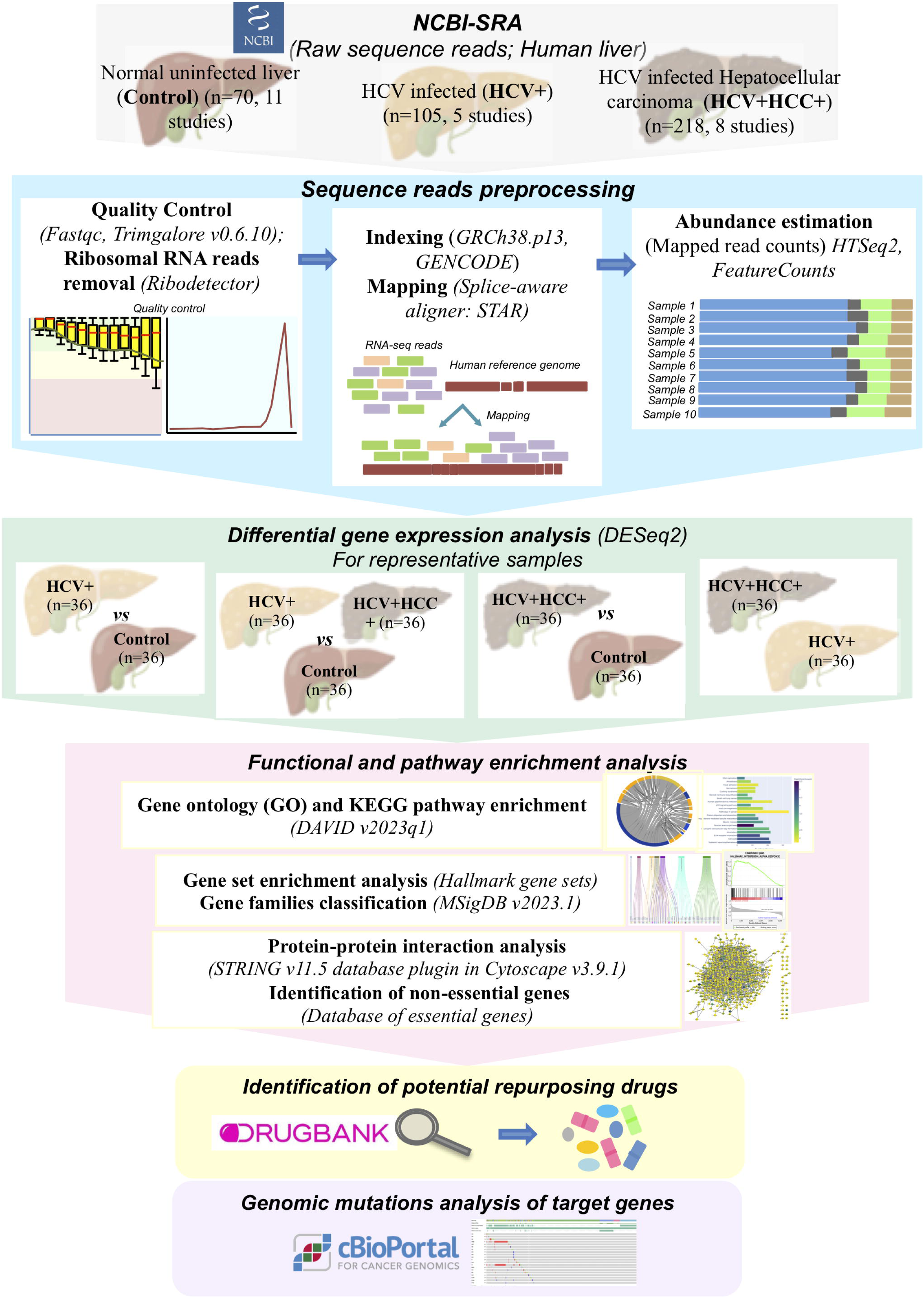
Overall workflow of this study to perform RNA sequencing data analysis on three types of human liver tissues: Hepatitis C virus (HCV) infected (HCV+), HCV induced/ infected hepatocellular carcinoma (HCV+HCC+) and normal non-infected (Control) available in NCBI-SRA. Differentially expressed genes were identified for four groups: (HCV+ vs Control), (HCV+HCC+ vs Control), (HCV+ AND HCV+HCC+ vs Control), and (HCV+HCC+ vs HCV+). Significant upregulated genes subjected to functional and pathway enrichment analyses, identification of non-essential genes. Identified non-essential genes were looked into the DrugBank repository for available drugs to identify potential repurposing drugs candidates. Genomic alterations were also looked into for these selected target genes.

### RNA-sequencing data collection

We searched the RNA-seq data for Hepatitis C virus (HCV) in NCBI-SRA using the keywords: ((Hepatitis C virus) OR HCV) AND “Homo sapiens” [orgn: _txid9606]. We found 6090 samples from this search. Further, we filtered these samples and selected human liver tissue samples from HCV infected patients (HCV+), and patients with HCV infection and HCC as primary disease (HCV+HCC+). We also looked for RNA-seq data of non-infected normal human liver samples (Control). We used human liver tissue samples for analysis as it is primarily affected during HCV infection and HCC condition. We have taken RNA-seq data for three types of human liver tissue samples: i) Healthy/normal liver, ii) HCV infected, and iii) HCV-related HCC and labelled them as Control, HCV+, HCV+HCC+, respectively. We retrieved 70, 105, and 218 samples from 11, 05, and 08 studies for Control, HCV+, and HCV+HCC+ categories, respectively (393 samples from 20 studies). We selected representative samples from three different categories based on read quality and mapped coverage within a particular study. To avoid biasness towards particular study or cohort, we included samples from all studies while reducing samples from studied having larger datasets. The representative samples from three different categories (36 for each category, 108 in total, from 20 studies) used in this study with their detailed information are given in **Supplementary Table S1**.

### Read quality assessment and rRNA removal

The selected RNA-seq read files were subjected to read quality assessment. We checked the quality of sequence reads by using Fastqc software (http://www.bioinformatics.babraham.ac.uk/projects/fastqc/). The sequencing adapter content was removed using Trimgalore v0.6.10. (https://github.com/FelixKrueger/TrimGalore). The ribosomal RNA reads were removed by using the program Ribodetector v0.2.7 using default parameters ^70^.

### Indexing and mapping to human genome

The reference human genome and its annotation [Release 43 (GRCh38.p13)] were retrieved from the GENCODE project ^71^ and used to generate indexes. We mapped the quality checked sequence reads to reference genome from indexed genome by using splice aware aligner STAR v2.7.10b ^72^. The mapped reads were quantified using HTseq-count v2.0.2 ^73^ and featureCounts v2.0.3 ^74^. The mapping and read counts outputs were summarised using MultiQC v1.13 ^75^.

### Differential gene expression analysis

The un-normalised mapped reads were subjected to counts normalization and differential gene expression analysis. We used DeSeq2, an R based Bioconductor package, which estimates scaling factors, gene-wise dispersion, and fits a generalized linear model for testing and generating a list of DEGs ^76^. We found DEGs in HCV+ samples (HCV+ vs Control), HCV+HCC+ (HCV+HCC+ vs Control), combining samples from both HCV+ and HCV+HCC+ (HCV+ AND HCV+HCC+ vs Control), and comparing HCV+HCC+ with HCV+ (HCV+HCC+ vs HCV+). Collectively, we analysed 324 samples in four groups: i) 72 (36 HCV+ vs 36 Control); ii) 72 (36 HCV+HCC+ vs 36 Control); iii) 108 (36 HCV+ AND 36 HCV+HCC+ vs 36 Control) and iv) 72 (36 HCV+HCC+ vs 36 HCV+). DEGs showing log2 fold change (log2FC) +≥ 1.5 and -≤ 1.5 with p-adjusted value <0.01 were selected as significant DEGs.

### Visualization of DEGs

The normalized counts and DEGs were visualized by dispersion estimation and mean-average (MA) plot using in-built functions ‘plotDispEsts’ and ‘plotMA’ respectively from the DeSeq2 package. We also generated volcano plots to get an overview of DEGs based on log2FC and p-adjusted values using an R based ‘enhancedvolcano’ package (https://github.com/kevinblighe/EnhancedVolcano).

### Functional and pathway enrichment analysis

We carried out GO and KEGG pathway enrichment analyses on the significantly upregulated genes for four study groups: (HCV+ vs Control), (HCV+HCC+ vs Control), (HCV+ AND HCV+HCC+ vs Control), and (HCV+HCC+ vs HCV+). We performed GO and KEGG enrichment analyses using the DAVID software with p-value cut-off of 0.05 ^77^. Top 20 GO annotations for each BP, CC and MF were visualized using the web server MonaGO ^78^. Top 20 enriched KEGG pathways are visualized in the form of bar plots using Python Plotly library.

### Gene set enrichment analysis

We identified the major gene families and top functional categories of DEGs from four study groups. For this, we carried out gene set enrichment analysis (GSEA). We used hallmark gene sets from Molecular Signature Database v2023.1 (MsigDB) ^79^ in GSEA software ^80^ for GSEA.

### Protein-protein interaction analysis

To study the interactions among the upregulated genes, we performed protein-protein interaction analysis. We searched for interactions among targeted upregulated genes using the STRING v11.5 database plugin in Cytoscape v3.9.1 using default parameters ^81, 82^. The network was visualized in Cytoscape v3.9.1, where each gene node was colored based on log2FC.

### Identification of essential and non-essential genes from DEGs

Essential genes are crucial for the survival and maintenance of an organism. We used the list of essential genes for *Homo sapiens* from the database of essential genes ^83^ for identification of essential genes. The identified upregulated genes were classified into essential and non-essential genes.

### Prediction of repurposing drugs targeting upregulated DEGs

The non-essential upregulated genes from four study groups were used as targets to look for the FDA-approved drugs from ‘DrugBank’ database v 5.1.10 ^84^. We also collected information about drugs such as drug group, drug type, and drug action. We also searched the literature for experimental, preclinical or clinical studies of these drugs for activity against HCV infection and HCC condition. We further prioritized the drugs as potential repurposing drug candidates fulfilling the criteria: i) drugs belonging to the drug groups-approved; approved, experimental; approved, investigational; approved, vet-approved, ii) drugs having action against target genes – antagonist, inhibitor, antibody, blocker, modulator, iii) not previously tested against HCV infection or/and HCC condition.

### Mutational profiling of prioritised target genes for drug repurposing

The genomic mutations were assessed for the identified non-essential genes from four studied groups. We used all the hepatocellular carcinoma related studies available in the cBioPortal web server (https://www.cbioportal.org/) for obtaining the mutational profiles of the tentative target genes.

### siRNAs and sgRNAs prediction for prioritised target genes

We also predicted small interfering RNAs (siRNAs) and small guide RNAs (sgRNAs) for Clustered regularly interspaced short palindromic repeats (CRISPR) for the prioritized target genes. We predicted the 19-mer siRNAs using support vector machine based binary predictive algorithm developed on homogenous dataset in the siRNApred web server (https://webs.iiitd.edu.in/raghava/sirnapred/). For sgRNA designing, we used CRISPick web server (https://portals.broadinstitute.org/gppx/crispick/public) for CRISPR interference (CRISPRi) using Human GRCh38(NCBI RefSeq v.GCF_000001405.40-RS_2023_10) for the enzyme SpyoCas9 ^85^.

## Abbreviations

BP: Biological processes
CC: Cellular components
DAAs: Direct-acting antivirals
DAVID: Database for annotation, visualization, and integrated discovery
DEGs: Differentially expressed genes
FDA: Food and drug administration
GO: Gene ontology
GSEA: Gene set enrichment analysis
HCC: Hepatocellular carcinoma
HCV: Hepatitis C virus
HCV+: HCV infected
HCV+HCC+: HCV related or induced HCC
HTAs: Host targeting agents
KEGG: Kyoto encyclopedia of genes and genomes
lfcSE: Standard error of log2 fold change
log2FC: Log2 fold change
MA: Mean-Average
MF: Molecular functions
MSigDB: Molecular signature database
NCBI-SRA: National center for biotechnology information-Sequence read archive
padj: Adjusted p-value
PPI: Protein-protein interaction
siRNAs: small interfering RNAs
sgRNAs: small guide RNAs
CRISPR: Clustered regularly interspaced short palindromic repeats
CRISPRi: CRISPR interference
SARS-CoV-2: severe acute respiratory syndrome coronavirus 2

## CRediT authorship contributions

**Sakshi Kamboj:** Methodology, Data curation, Formal analysis, Investigation, Validation, Visualization, Writing-Original draft, Writing-Review & Editing

**Manoj Kumar:** Conceptualization, Supervision, Validation, Funding acquisition, Investigation, Writing-review & editing

## Funding

We acknowledge the support from Council of Scientific and Industrial Research (CSIR) - Institute of Microbial Technology grant (project no.: OLP0192) and CSIR - Senior Research Fellowship (CSIR-SRF), Award No. 31/050(0453)/2019-EMR-I to Sakshi Kamboj.

## Conflict of interest statement

The authors declare that they have no competing interests.

## Supplementary data availability

**Supplementary file 1.** A document file containing detailed methodology and all Supplementary Figures S1-S17.

**Supplementary file 2.** An excel file containing Supplementary Tables S1-S3.

**Supplementary file 3.** An excel file containing Supplementary Tables S4-S23.

## Notes

### Competing Interest Statement

The authors have declared no competing interest.

## References

1. Villanueva A. Hepatocellular Carcinoma. N Engl J Med. Apr 11 2019;380(15):1450–1462. doi:10.1056/NEJMra1713263

2. Llovet JM, Montal R, Sia D, Finn RS. Molecular therapies and precision medicine for hepatocellular carcinoma. Nat Rev Clin Oncol. Oct 2018;15(10):599–616. doi:10.1038/s41571-018-0073-4

3. Llovet JM, Kelley RK, Villanueva A, et al. Hepatocellular carcinoma. Nat Rev Dis Primers. Jan 21 2021;7(1):6. doi:10.1038/s41572-020-00240-3

4. Llovet JM, Pinyol R, Kelley RK, et al. Molecular pathogenesis and systemic therapies for hepatocellular carcinoma. Nat Cancer. Apr 2022;3(4):386–401. doi:10.1038/s43018-022-00357-2

5. Moradpour D, Penin F, Rice CM. Replication of hepatitis C virus. Nat Rev Microbiol. Jun 2007;5(6):453–63. doi:10.1038/nrmicro1645

6. Lingala S, Ghany MG. Natural History of Hepatitis C. Gastroenterol Clin North Am. Dec 2015;44(4):717–34. doi:10.1016/j.gtc.2015.07.003

7. Pawlotsky JM. Treatment failure and resistance with direct-acting antiviral drugs against hepatitis C virus. Hepatology. May 2011;53(5):1742–51. doi:10.1002/hep.24262

8. Crouchet E, Wrensch F, Schuster C, Zeisel MB, Baumert TF. Host-targeting therapies for hepatitis C virus infection: current developments and future applications. Therap Adv Gastroenterol. 2018;11:1756284818759483. doi:10.1177/1756284818759483

9. Hu L, Li J, Cai H, et al. Avasimibe: A novel hepatitis C virus inhibitor that targets the assembly of infectious viral particles. Antiviral Res. Dec 2017;148:5–14. doi:10.1016/j.antiviral.2017.10.016

10. Bobardt M, Ramirez CM, Baum MM, Ure D, Foster R, Gallay PA. The combination of the NS5A and cyclophilin inhibitors results in an additive anti-HCV inhibition in humanized mice without development of resistance. PLoS One. 2021;16(5):e0251934. doi:10.1371/journal.pone.0251934

11. Huang A, Yang XR, Chung WY, Dennison AR, Zhou J. Targeted therapy for hepatocellular carcinoma. Signal Transduct Target Ther. Aug 11 2020;5(1):146. doi:10.1038/s41392-020-00264-x

12. Thakur A, Kumar M. Integration of Human and Viral miRNAs in Epstein-Barr Virus-Associated Tumors and Implications for Drug Repurposing. Omics. Mar 2023;27(3):93–108. doi:10.1089/omi.2023.0005

13. Zhou A, Dong X, Liu M, Tang B. Comprehensive Transcriptomic Analysis Identifies Novel Antiviral Factors Against Influenza A Virus Infection. Front Immunol. 2021;12:632798. doi:10.3389/fimmu.2021.632798

14. Hoshida Y, Villanueva A, Sangiovanni A, et al. Prognostic gene expression signature for patients with hepatitis C-related early-stage cirrhosis. Gastroenterology. May 2013;144(5):1024–30. doi:10.1053/j.gastro.2013.01.021

15. Cheng J, Chen Z, Zuo G, Cao W. Integrated analysis of differentially expressed genes, differentially methylated genes, and natural compounds in hepatitis C virus-induced cirrhosis. J Int Med Res. Jan 2022;50(1):3000605221074525. doi:10.1177/03000605221074525

16. Li YC, Hu WY, Li CH, et al. Differential expression and significance of 5-hydroxymethylcytosine modification in hepatitis B virus carriers and patients with liver cirrhosis and liver cancer. World J Gastrointest Surg. Mar 27 2023;15(3):346–361. doi:10.4240/wjgs.v15.i3.346

17. Bièche I, Asselah T, Laurendeau I, et al. Molecular profiling of early stage liver fibrosis in patients with chronic hepatitis C virus infection. Virology. Feb 5 2005;332(1):130–44. doi:10.1016/j.virol.2004.11.009

18. Wróblewska A, Bernat A, Woziwodzka A, et al. Interferon lambda polymorphisms associate with body iron indices and hepatic expression of interferon-responsive long non-coding RNA in chronic hepatitis C. Clin Exp Med. May 2017;17(2):225–232. doi:10.1007/s10238-016-0423-4

19. Ma J, Bai H, Gong T, et al. Novel skewed usage of B-cell receptors in COVID-19 patients with various clinical presentations. Immunol Lett. Sep 2022;249:23–32. doi:10.1016/j.imlet.2022.08.006

20. Wood O, Woo J, Seumois G, et al. Gene expression analysis of TIL rich HPV-driven head and neck tumors reveals a distinct B-cell signature when compared to HPV independent tumors. Oncotarget. Aug 30 2016;7(35):56781–56797. doi:10.18632/oncotarget.10788

21. Yao C, Xu R, Li Q, et al. Identification and validation of an immunological microenvironment signature and prediction model for epstein-barr virus positive lymphoma: Implications for immunotherapy. Front Oncol. 2022;12:970544. doi:10.3389/fonc.2022.970544

22. Horsmans Y, Berg T, Desager JP, et al. Isatoribine, an agonist of TLR7, reduces plasma virus concentration in chronic hepatitis C infection. Hepatology. Sep 2005;42(3):724–31. doi:10.1002/hep.20839

23. Pockros PJ, Guyader D, Patton H, et al. Oral resiquimod in chronic HCV infection: safety and efficacy in 2 placebo-controlled, double-blind phase IIa studies. J Hepatol. Aug 2007;47(2):174–82. doi:10.1016/j.jhep.2007.02.025

24. Lowe HI, Toyang NJ, McLaughlin W. Potential of Cannabidiol for the Treatment of Viral Hepatitis. Pharmacognosy Res. Jan-Mar 2017;9(1):116–118. doi:10.4103/0974-8490.199780

25. Liu Z, Pu Y, Bao Y, He S. Investigation of Potential Molecular Biomarkers for Diagnosis and Prognosis of AFP-Negative HCC. Int J Gen Med. 2021;14:4369–4380. doi:10.2147/ijgm.S323868

26. Ferro I, Gavini J, Gallo S, et al. The human vault RNA enhances tumorigenesis and chemoresistance through the lysosome in hepatocellular carcinoma. Autophagy. Jan 2022;18(1):191–203. doi:10.1080/15548627.2021.1922983

27. Wu W, Li Q, Zhu Z, et al. HTR1D functions as a key target of HOXA10-AS/miR-340-3p axis to promote the malignant outcome of pancreatic cancer via PI3K-AKT signaling pathway. Int J Biol Sci. 2022;18(9):3777–3794. doi:10.7150/ijbs.70546

28. Sigala S, Bothou C, Penton D, et al. A Comprehensive Investigation of Steroidogenic Signaling in Classical and New Experimental Cell Models of Adrenocortical Carcinoma. Cells. Apr 24 2022;11(9)doi:10.3390/cells11091439

29. Neubauer E, Latif M, Krause J, et al. Up regulation of the steroid hormone synthesis regulator HSD3B2 is linked to early PSA recurrence in prostate cancer. Exp Mol Pathol. Aug 2018;105(1):50–56. doi:10.1016/j.yexmp.2018.05.006

30. Lin JC, Liu CL, Chang YC, et al. Trilostane, a 3β-hydroxysteroid dehydrogenase inhibitor, suppresses growth of hepatocellular carcinoma and enhances anti-cancer effects of sorafenib. Invest New Drugs. Dec 2021;39(6):1493–1506. doi:10.1007/s10637-021-01132-3

31. Maurya V, Kumar P, Chakraborti S, et al. Zolmitriptan attenuates hepatocellular carcinoma via activation of caspase mediated apoptosis. Chem Biol Interact. Aug 1 2019;308:120–129. doi:10.1016/j.cbi.2019.05.033

32. Cheng X, Wu H, Jin ZJ, et al. Up-regulation of chemokine receptor CCR4 is associated with Human Hepatocellular Carcinoma malignant behavior. Sci Rep. Sep 28 2017;7(1):12362. doi:10.1038/s41598-017-10267-4

33. Fu SJ, Shen SL, Li SQ, et al. Hornerin promotes tumor progression and is associated with poor prognosis in hepatocellular carcinoma. BMC Cancer. Aug 13 2018;18(1):815. doi:10.1186/s12885-018-4719-5

34. Shimizu Y, Mizuno S, Fujinami N, et al. Plasma and tumoral glypican-3 levels are correlated in patients with hepatitis C virus-related hepatocellular carcinoma. Cancer Sci. Feb 2020;111(2):334–342. doi:10.1111/cas.14251

35. Sixtos-Alonso MS, Sánchez-Muñoz F, Sánchez-Ávila JF, et al. IFN-stimulated gene expression is a useful potential molecular marker of response to antiviral treatment with Peg-IFNα 2b and ribavirin in patients with hepatitis C virus genotype 1. Arch Med Res. Jan 2011;42(1):28–33. doi:10.1016/j.arcmed.2011.01.001

36. Wu J, Qu J, Cao H, et al. Monoclonal antibody AC10364 inhibits cell proliferation in 5-fluorouracil resistant hepatocellular carcinoma via apoptotic pathways. Onco Targets Ther. 2019;12:5053–5067. doi:10.2147/ott.S206517

37. Wong LR, Zheng J, Wilhelmsen K, et al. Eicosanoid signalling blockade protects middle-aged mice from severe COVID-19. Nature. May 2022;605(7908):146–151. doi:10.1038/s41586-022-04630-3

38. Kim JH, Lee ES, Yun J, et al. Calsequestrin 2 overexpression in breast cancer increases tumorigenesis and metastasis by modulating the tumor microenvironment. Mol Oncol. Jan 2022;16(2):466–484. doi:10.1002/1878-0261.13136

39. Abedini F, Amjadi O, Hedayatizadeh-Omran A, Lira SA, Ahangari G. Serotonin Receptors and Acetylcholinesterase Gene Expression Alternations: The Potential Value on Tumor Microenvironment of Gastric Cancer. Oncology. 2023;101(7):415–424. doi:10.1159/000530878

40. Sangro B, Gomez-Martin C, de la Mata M, et al. A clinical trial of CTLA-4 blockade with tremelimumab in patients with hepatocellular carcinoma and chronic hepatitis C. J Hepatol. Jul 2013;59(1):81–8. doi:10.1016/j.jhep.2013.02.022

41. Ren Z, Ducreux M, Abou-Alfa GK, et al. Tislelizumab in Patients with Previously Treated Advanced Hepatocellular Carcinoma (RATIONALE-208): A Multicenter, Non-Randomized, Open-Label, Phase 2 Trial. Liver Cancer. Feb 2023;12(1):72–84. doi:10.1159/000527175

42. Ozunal ZG, Cakil YD, Isan H, Saglam E, Aktas RG. Sertraline in combination with sorafenib: A promising pharmacotherapy to target both depressive disorders and hepatocellular cancer. Biol Futur. Dec 2019;70(4):341–348. doi:10.1556/019.70.2019.39

43. Liu G, Sun BY, Sun J, et al. BRG1 regulates lipid metabolism in hepatocellular carcinoma through the PIK3AP1/PI3K/AKT pathway by mediating GLMP expression. Dig Liver Dis. May 2022;54(5):692–700. doi:10.1016/j.dld.2021.05.002

44. Gu X, Mao Y, Shi C, et al. MAGEC2 Correlates With Unfavorable Prognosis And Promotes Tumor Development In HCC Via Epithelial-Mesenchymal Transition. Onco Targets Ther. 2019;12:7843–7855. doi:10.2147/ott.S213164

45. Lu JG, Nguyen L, Samadzadeh S, et al. Expression of proteins upregulated in hepatocellular carcinoma in patients with alcoholic hepatitis (AH) compared to non-alcoholic steatohepatitis (NASH): An immunohistochemical analysis of candidate proteins. Exp Mol Pathol. Apr 2018;104(2):125–129. doi:10.1016/j.yexmp.2018.02.001

46. Lu SX, Zhang CZ, Luo RZ, et al. Zic2 promotes tumor growth and metastasis via PAK4 in hepatocellular carcinoma. Cancer Lett. Aug 28 2017;402:71–80. doi:10.1016/j.canlet.2017.05.018

47. Zhang MH, Niu H, Li Z, Huo RT, Wang JM, Liu J. Activation of PI3K/AKT is involved in TINAG-mediated promotion of proliferation, invasion and migration of hepatocellular carcinoma. Cancer Biomark. 2018;23(1):33–43. doi:10.3233/cbm-181277

48. Molania R, Mahjoubi F, Mirzaei R, Khatami SR, Mahjoubi B. A Panel of Cancer Testis Antigens and Clinical Risk Factors to Predict Metastasis in Colorectal Cancer. J Biomark. 2014;2014:272683. doi:10.1155/2014/272683

49. Gong J, Tian J, Lou J, et al. A polymorphic MYC response element in KBTBD11 influences colorectal cancer risk, especially in interaction with an MYC-regulated SNP rs6983267. Ann Oncol. Mar 1 2018;29(3):632–639. doi:10.1093/annonc/mdx789

50. Li MW, Huang FX, Xie ZC, Hong HY, Xu QY, Peng ZG. Identification of three small nucleolar RNAs (snoRNAs) as potential prognostic markers in diffuse large B-cell lymphoma. Cancer Med. Feb 2023;12(3):3812–3829. doi:10.1002/cam4.5115

51. Xie ZS, Han XY, Zhou ZY, et al. Design and synthesis of dabigatran etexilate derivatives with inhibiting thrombin activity for hepatocellular carcinoma treatment. Biomed Pharmacother. Jan 2024;170:116018. doi:10.1016/j.biopha.2023.116018

52. Guo P, Hu Q, Wang J, Hai L, Nie X, Zhao Q. Butorphanol inhibits angiogenesis and migration of hepatocellular carcinoma and regulates MAPK pathway. J Antibiot (Tokyo). Nov 2022;75(11):626–634. doi:10.1038/s41429-022-00565-z

53. Neumann-Haefelin C, Thimme R. Adaptive immune responses in hepatitis C virus infection. Curr Top Microbiol Immunol. 2013;369:243–62. doi:10.1007/978-3-642-27340-7_10

54. Reungoat E, Grigorov B, Zoulim F, Pécheur EI. Molecular Crosstalk between the Hepatitis C Virus and the Extracellular Matrix in Liver Fibrogenesis and Early Carcinogenesis. Cancers (Basel). May 9 2021;13(9)doi:10.3390/cancers13092270

55. Sepulveda-Crespo D, Resino S, Martinez I. Strategies Targeting the Innate Immune Response for the Treatment of Hepatitis C Virus-Associated Liver Fibrosis. Drugs. Mar 2021;81(4):419–443. doi:10.1007/s40265-020-01458-x

56. Peña-Asensio J, Calvo H, Torralba M, Miquel J, Sanz-de-Villalobos E, Larrubia JR. Gamma-Chain Receptor Cytokines & PD-1 Manipulation to Restore HCV-Specific CD8(+) T Cell Response during Chronic Hepatitis C. Cells. Mar 3 2021;10(3)doi:10.3390/cells10030538

57. Chen QT, Zhang ZY, Huang QL, et al. HK1 from hepatic stellate cell-derived extracellular vesicles promotes progression of hepatocellular carcinoma. Nat Metab. Oct 2022;4(10):1306–1321. doi:10.1038/s42255-022-00642-5

58. Nimitrungtawee N, Inmutto N, Chattipakorn SC, Chattipakorn N. Extracellular vesicles as a new hope for diagnosis and therapeutic intervention for hepatocellular carcinoma. Cancer Med. Dec 2021;10(23):8253–8271. doi:10.1002/cam4.4370

59. Wong BHS, Poh ZS, Wei JTC, et al. High Extracellular K(+) Skews T-Cell Differentiation Towards Tumour Promoting Th2 and T(reg) Subsets. Eur J Immunol. Dec 9 2024:e202451440. doi:10.1002/eji.202451440

60. Bergqvist A, Sundström S, Dimberg LY, Gylfe E, Masucci MG. The hepatitis C virus core protein modulates T cell responses by inducing spontaneous and altering T-cell receptor-triggered Ca2+ oscillations. J Biol Chem. May 23 2003;278(21):18877–83. doi:10.1074/jbc.M300185200

61. Lima LDP, Machado CJ, Rodrigues J, et al. Immunohistochemical Coexpression of Epithelial Cell Adhesion Molecule and Alpha-Fetoprotein in Hepatocellular Carcinoma. Can J Gastroenterol Hepatol. 2018;2018:5970852. doi:10.1155/2018/5970852

62. Sanchez JI, Jiao J, Kwan SY, et al. Lipidomic Profiles of Plasma Exosomes Identify Candidate Biomarkers for Early Detection of Hepatocellular Carcinoma in Patients with Cirrhosis. Cancer Prev Res (Phila). Oct 2021;14(10):955–962. doi:10.1158/1940-6207.Capr-20-0612

63. Zhou L, Zhang Q, Deng H, Ou S, Liang T, Zhou J. The SNHG1-Centered ceRNA Network Regulates Cell Cycle and Is a Potential Prognostic Biomarker for Hepatocellular Carcinoma. Tohoku J Exp Med. Nov 11 2022;258(4):265–276. doi:10.1620/tjem.2022.J083

64. Perlemuter G, Cacoub P, Sbaï A, et al. Hepatitis C virus infection in systemic lupus erythematosus: a case-control study. J Rheumatol. Jul 2003;30(7):1473–8.

65. Vucur M, Ghallab A, Schneider AT, et al. Sublethal necroptosis signaling promotes inflammation and liver cancer. Immunity. Jul 11 2023;56(7):1578–1595.e8. doi:10.1016/j.immuni.2023.05.017

66. Broering R, Trippler M, Werner M, et al. Hepatic expression of proteasome subunit alpha type-6 is upregulated during viral hepatitis and putatively regulates the expression of ISG15 ubiquitin-like modifier, a proviral host gene in hepatitis C virus infection. J Viral Hepat. May 2016;23(5):375–86. doi:10.1111/jvh.12508

67. White IR, Patel K, Symonds WT, et al. Serum proteomic analysis focused on fibrosis in patients with hepatitis C virus infection. J Transl Med. Jul 11 2007;5:33. doi:10.1186/1479-5876-5-33

68. Odigie M, Osinusi A, Barrett L, et al. Inteleukin-23 promotes interferon-α responsiveness in hepatitis C virus/HIV-coinfected patients. AIDS Res Hum Retroviruses. Aug 2014;30(8):775–82. doi:10.1089/aid.2014.0003

69. Dillon ST, Bhasin MK, Feng X, Koh DW, Daoud SS. Quantitative proteomic analysis in HCV-induced HCC reveals sets of proteins with potential significance for racial disparity. J Transl Med. Oct 1 2013;11:239. doi:10.1186/1479-5876-11-239

70. Deng ZL, Münch PC, Mreches R, McHardy AC. Rapid and accurate identification of ribosomal RNA sequences via deep learning. Nucleic Acids Res. Jun 10 2022;50(10):e60. doi:10.1093/nar/gkac112

71. Frankish A, Diekhans M, Ferreira AM, et al. GENCODE reference annotation for the human and mouse genomes. Nucleic Acids Res. Jan 8 2019;47(D1):D766–d773. doi:10.1093/nar/gky955

72. Dobin A, Davis CA, Schlesinger F, et al. STAR: ultrafast universal RNA-seq aligner. Bioinformatics. Jan 1 2013;29(1):15–21. doi:10.1093/bioinformatics/bts635

73. Anders S, Pyl PT, Huber W. HTSeq--a Python framework to work with high-throughput sequencing data. Bioinformatics. Jan 15 2015;31(2):166–9. doi:10.1093/bioinformatics/btu638

74. Liao Y, Smyth GK, Shi W. featureCounts: an efficient general purpose program for assigning sequence reads to genomic features. Bioinformatics. Apr 1 2014;30(7):923–30. doi:10.1093/bioinformatics/btt656

75. Ewels P, Magnusson M, Lundin S, Käller M. MultiQC: summarize analysis results for multiple tools and samples in a single report. Bioinformatics. Oct 1 2016;32(19):3047–8. doi:10.1093/bioinformatics/btw354

76. Love MI, Huber W, Anders S. Moderated estimation of fold change and dispersion for RNA-seq data with DESeq2. Genome Biol. 2014;15(12):550. doi:10.1186/s13059-014-0550-8

77. Sherman BT, Hao M, Qiu J, et al. DAVID: a web server for functional enrichment analysis and functional annotation of gene lists (2021 update). Nucleic Acids Res. Jul 5 2022;50(W1):W216–w221. doi:10.1093/nar/gkac194

78. Xin Z, Cai Y, Dang LT, et al. MonaGO: a novel gene ontology enrichment analysis visualisation system. BMC Bioinformatics. Feb 14 2022;23(1):69. doi:10.1186/s12859-022-04594-1

79. Liberzon A, Birger C, Thorvaldsdóttir H, Ghandi M, Mesirov JP, Tamayo P. The Molecular Signatures Database (MSigDB) hallmark gene set collection. Cell Syst. Dec 23 2015;1(6):417–425. doi:10.1016/j.cels.2015.12.004

80. Subramanian A, Tamayo P, Mootha VK, et al. Gene set enrichment analysis: a knowledge-based approach for interpreting genome-wide expression profiles. Proc Natl Acad Sci U S A. Oct 25 2005;102(43):15545–50. doi:10.1073/pnas.0506580102

81. Szklarczyk D, Gable AL, Nastou KC, et al. The STRING database in 2021: customizable protein-protein networks, and functional characterization of user-uploaded gene/measurement sets. Nucleic Acids Res. Jan 8 2021;49(D1):D605–d612. doi:10.1093/nar/gkaa1074

82. Shannon P, Markiel A, Ozier O, et al. Cytoscape: a software environment for integrated models of biomolecular interaction networks. Genome Res. Nov 2003;13(11):2498–504. doi:10.1101/gr.1239303

83. Luo H, Lin Y, Liu T, et al. DEG 15, an update of the Database of Essential Genes that includes built-in analysis tools. Nucleic Acids Res. Jan 8 2021;49(D1):D677–d686. doi:10.1093/nar/gkaa917

84. Wishart DS, Feunang YD, Guo AC, et al. DrugBank 5.0: a major update to the DrugBank database for 2018. Nucleic Acids Res. Jan 4 2018;46(D1):D1074–d1082. doi:10.1093/nar/gkx1037

85. Sanson KR, Hanna RE, Hegde M, et al. Optimized libraries for CRISPR-Cas9 genetic screens with multiple modalities. Nat Commun. Dec 21 2018;9(1):5416. doi:10.1038/s41467-018-07901-8

