## Supplementary File 1 for "Integrated transcriptomic analyses identifies host-targeting repurposing drugs for hepatitis C virus infection and related hepatocellular carcinoma"

This document file includes **detailed methodology** and **Supplementary Figures** **S1** to **S17**.

**Other Supplementary Materials for this manuscript include the following:**

Supplementary file 2 containing **Supplementary Table S1** to **S3** (included in excel spreadsheets)

Supplementary file 3 containing **Supplementary Table S4** to **S23** (included in excel spreadsheets).

**Detailed Methodology**

**RNA-sequencing data collection:** We searched the RNA-seq data for Hepatitis C virus (HCV) in NCBI-SRA using the keywords: ((Hepatitis C virus) OR HCV) AND “Homo sapiens” [orgn: _txid9606]. We found 6090 samples from this search. Later on, we filtered these samples and selected human liver tissue samples from HCV infected patients (HCV+), and patients with HCV infection and HCC as primary disease (HCV+HCC+). We also looked for RNA-seq data of non-infected normal human liver samples (Control). We selected HCV infected patients samples from the studies: PRJEB27201 (55), PRJNA328986 (40), PRJNA485359 (6), PRJNA533227 (1) and PRJNA506130 (3). However, for HCV-infected HCC patients (HCV+HCC+), we utilized data from the studies: PRJDB10863 (2), PRJNA324260 (8), PRJNA591153 (5), PRJNA592558 (99), PRJNA592565 (23), PRJNA793464 (5), PRJNA770529 (10), and PRJNA928394 (66).

For control samples, we utilized representative control liver samples from nine studies: PRJNA328986 (6), PRJNA591153 (5), PRJNA30709 (3), PRJNA523510 (14), PRJNA558821 (9), PRJNA629495 (9), PRJNA683913 (3), PRJNA726931 (4), PRJNA324260 (10), PRJNA770529 (4), and PRJNA764684 (3). We used human liver tissue samples for analysis as it is primarily affected during HCV infection and HCC condition. We have taken RNA-seq data for three types of human liver tissue samples: i) Healthy/normal liver, ii) HCV infected, and iii) HCV-related HCC and labelled them as Control, HCV+, HCV+HCC+, respectively. We retrieved 70, 105, and 218 samples from 11, 05, and 08 studies for Control, HCV+, and HCV+HCC+ categories, respectively (393 samples from 20 studies). We selected representative samples from three different categories based on read quality and mapped coverage within a particular study. To avoid biasness towards particular study or cohort, we included samples from all studies while reducing samples from studied having larger datasets. The representative samples from three different categories (36 for each category, 108 in total, from 20 studies) used in this study are given in **Table 2**. The information on representative SRA runs employed in our study is provided in **Supplementary Table S1**.

**Read quality assessment and rRNA removal:** The selected RNA-seq read files were subjected to read quality assessment. We checked the quality of sequence reads by using Fastqc software (http://www.bioinformatics.babraham.ac.uk/projects/fastqc/). The sequencing adapter content was removed using Trimgalore v0.6.10. (https://github.com/FelixKrueger/TrimGalore). The ribosomal RNA reads were removed by using the program Ribodetector v0.2.7 using default parameters ^1^.

**Indexing and mapping to human genome:** The reference human genome and its annotation [Release 43 (GRCh38.p13)] were retrieved from the GENCODE project ^2^. The reference genome was processed to generate indexes of the genome. This indexing step is essential for mapping sequence reads to the reference genome. We mapped the quality checked sequence reads to reference genome from indexed genome by using splice aware aligner STAR v2.7.10b (Spliced transcripts alignment to a reference) using default parameters ^3^. The mapped reads were quantified using HTseq-count v2.0.2 using default parameters with intersection-nonempty overlapping mode ^4^and featureCounts v2.0.3 using default parameters ^5^. The mapping and read counts outputs were summarised using MultiQC v1.13 using default parameters ^6^.

**Differential gene expression analysis:** The un-normalised mapped reads were subjected to counts normalization and differential gene expression analysis. We used DeSeq2, an R based Bioconductor package, which estimates scaling factors, gene-wise dispersion, and fits a generalized linear model for testing and generating a list of DEGs ^7^. We found DEGs in HCV+ samples (HCV+ vs Control), HCV+HCC+ (HCV+HCC+ vs Control), combining samples from both HCV+ and HCV+HCC+ (HCV+ AND HCV+HCC+ vs Control), and comparing HCV+HCC+ with HCV+ (HCV+HCC+ vs HCV+). Collectively, we analysed 324 samples in four groups: i) 72 (36 HCV+ vs 36 Control); ii) 72 (36 HCV+HCC+ vs 36 Control); iii) 108 (36 HCV+ AND 36 HCV+HCC+ vs 36 Control) and iv) 72 (36 HCV+HCC+ vs 36 HCV+). DEGs showing log2 fold change (log2FC) +≥ 1.5 and -≤ 1.5 with p-adjusted value <0.01 were selected as significant DEGs.

**Visualization of DEGs:** The normalized counts and DEGs were visualized by dispersion estimation and mean-average (MA) plot using in-built functions ‘plotDispEsts’ and ‘plotMA’ respectively from the DeSeq2 package. We also generated volcano plots to get an overview of DEGs based on log2FC and p-adjusted values using an R based ‘enhancedvolcano’ package (https://github.com/kevinblighe/EnhancedVolcano).

**Functional and pathway enrichment analysis:** We carried out GO and KEGG pathway enrichment analyses on the significantly upregulated genes. We identified the important biological processes (BP), cellular components (CC), molecular functions (MF), and involved KEGG pathways for four study groups: (HCV+ vs Control), (HCV+HCC+ vs Control), (HCV+ AND HCV+HCC+ vs Control), and (HCV+HCC+ vs HCV+). We performed GO and KEGG enrichment analyses using the DAVID software with p-value cut-off of 0.05 ^8^. Top 20 GO annotations for each BP, CC and MF were visualized using the web server MonaGO ^9^. Top 20 enriched KEGG pathways are visualized in the form of bar plots using Python Plotly library.

**siRNAs and sgRNAs prediction for prioritised target genes:** We also predicted small interfering RNAs (siRNAs) and small guide RNAs (sgRNAs) for Clustered regularly interspaced short palindromic repeats (CRISPR) for the prioritized target genes. For siRNAs prediction, we downloaded the transcript sequences of all the target genes from NCBI GenBank. We predicted the 19-mer siRNAs using support vector machine based binary predictive algorithm developed on homogenous dataset in the siRNApred web server (<https://webs.iiitd.edu.in/raghava/sirnapred/>). For sgRNA designing, we used CRISPick web server (https://portals.broadinstitute.org/gppx/crispick/public) for CRISPR interference (CRISPRi) using Human GRCh38(NCBI RefSeq v.GCF_000001405.40-RS_2023_10) for the enzyme SpyoCas9 ^16^. This tool picks sgRNAs based on a constraint that it targets within 5-65% of the protein-coding region of target gene with on-target score of ≥ 0.2.

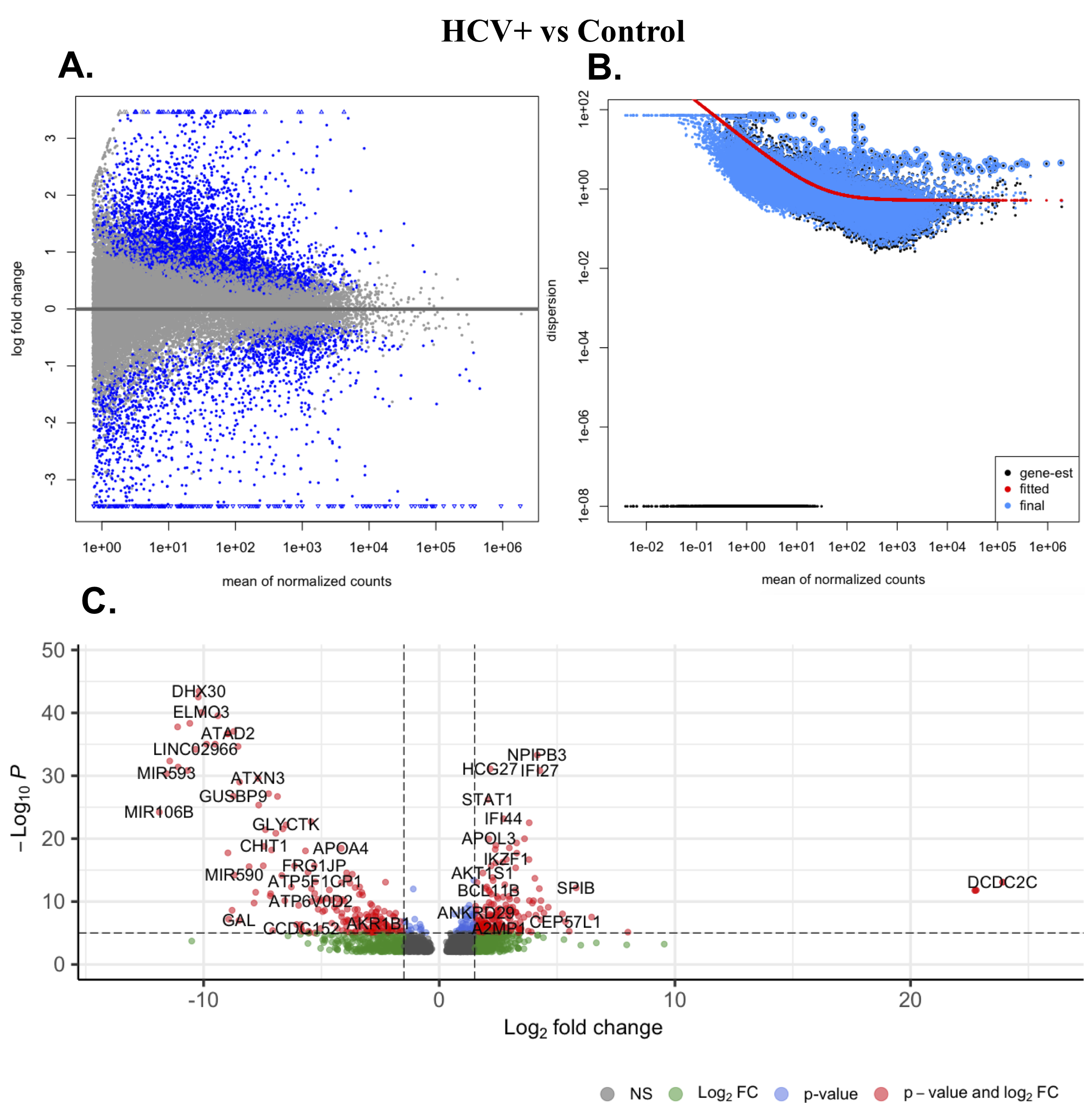

**Figure S1.** Visualizations of differential gene expression analysis of HCV+ vs Control **A.** MA plot shows the relationship between the normalized counts and log2FC at p-adjusted value <0.05 **B.** Dispersion plot showing the gene wise dispersion estimates and fitted estimates **C.** Volcano plot showing the significant DEGs with padj < 0.01 and log2FC cut-off 1.5.

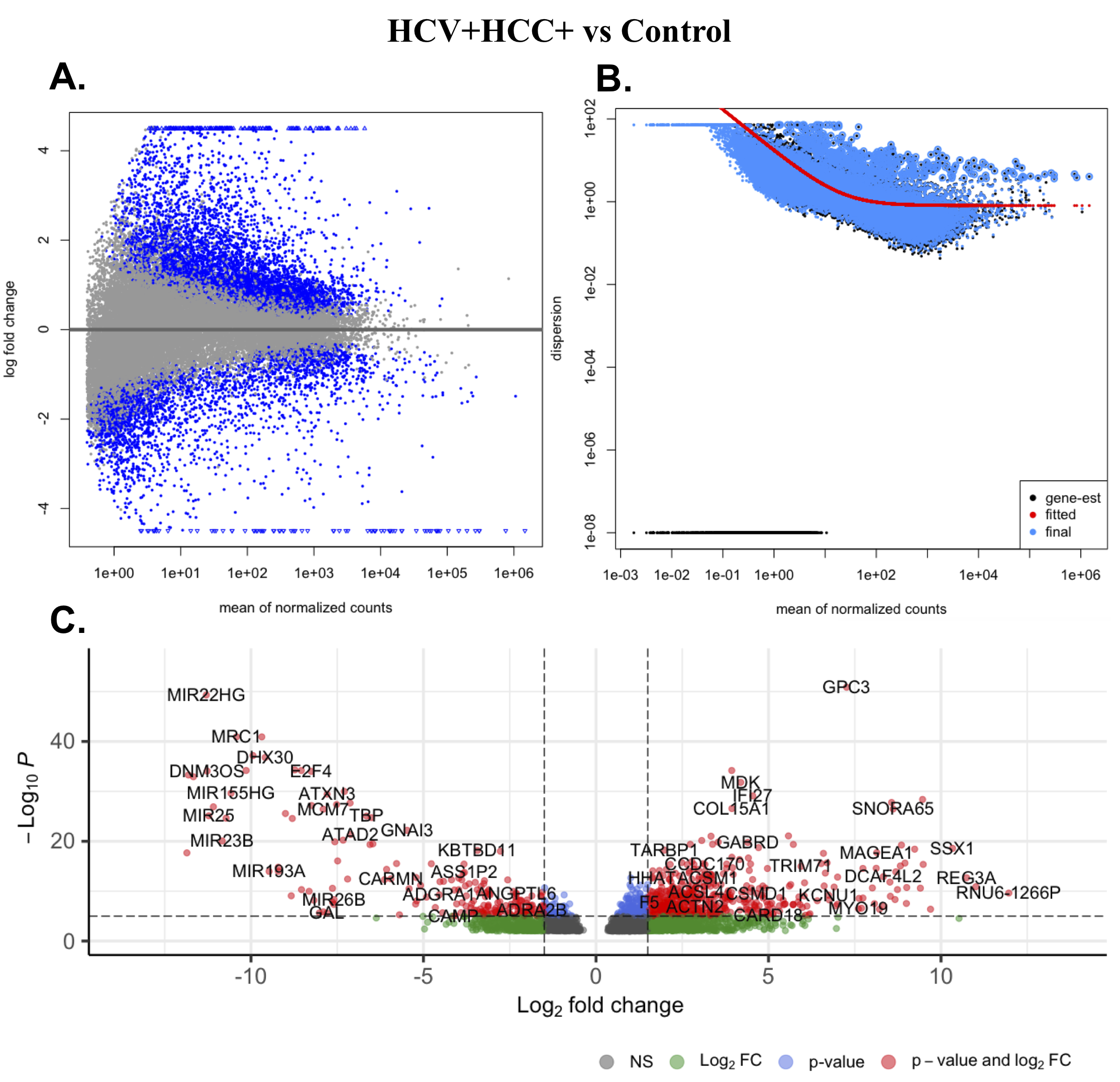

**Figure S2.** Visualizations of differential gene expression analysis of HCV+HCC+ vs Control **A.** MA plot shows the relationship between the normalized counts and log2FC at p-adjusted value <0.05 **B.** Dispersion plot showing the gene wise dispersion estimates and fitted estimates **C.** Volcano plot showing the significant DEGs with padj < 0.01 and log2FC cut-off 1.5.

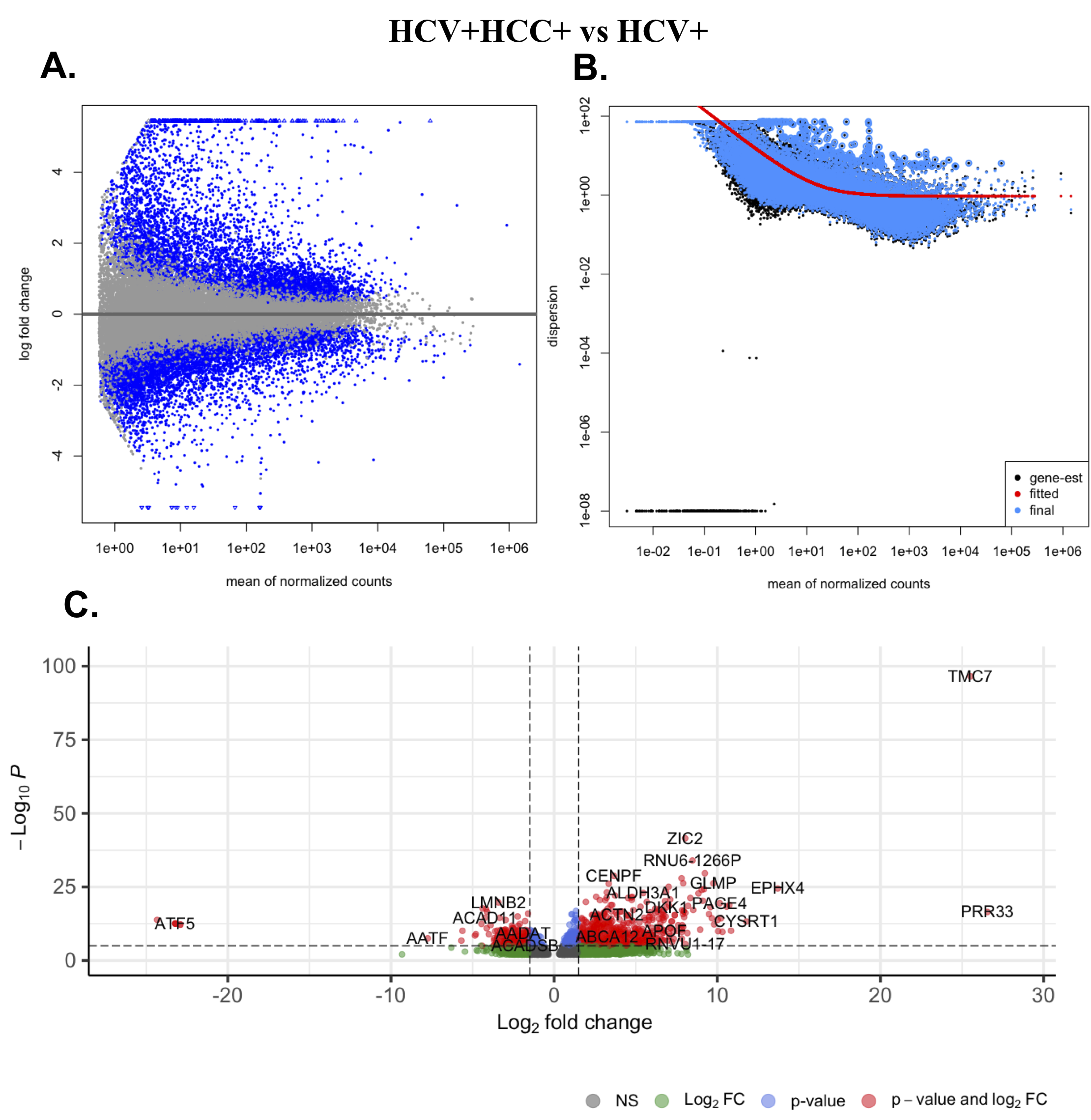

**Figure S3.** Visualizations of differential gene expression analysis of HCV+HCC+ vs HCV+ **A.** MA plot shows the relationship between the normalized counts and log2FC at p-adjusted value <0.05 **B.** Dispersion plot showing the gene wise dispersion estimates and fitted estimates **C.** Volcano plot showing the significant DEGs with padj < 0.01 and log2FC cut-off 1.5.

**Figure S4.** Gene Ontology and KEGG pathway enrichment for HCV+ vs Control group upregulated genes. Top 20 gene ontology terms in **A.** Biological processes **B.** Cellular component **C.** Molecular Functions. **D.** Top 20 enriched KEGG pathways. For A-C, the green arcs on the outside parallel to the main chord diagram denote possible (hierarchical) clusters. The number on an arc-node represents the percentage of common genes between two nodes/clusters. The grey links inside the chord diagram connect pairs of GO term/clusters, and indicates the existence of common genes between them.

**Figure S5.** Gene Ontology and KEGG pathway enrichment for HCV+HCC+ vs Control group upregulated genes. Top 20 gene ontology terms in **A.** Biological processes **B.** Cellular component **C.** Molecular Functions. **D.** Top 20 enriched KEGG pathways. For A-C, the green arcs on the outside parallel to the main chord diagram denote possible (hierarchical) clusters. The number on an arc-node represents the percentage of common genes between two nodes/clusters. The grey links inside the chord diagram connect pairs of GO term/clusters, and indicates the existence of common genes between them.

**Figure S6.** Gene Ontology and KEGG pathway enrichment for HCV+HCC+ vs HCV+ group upregulated genes. Top 20 gene ontology terms in **A.** Biological processes **B.** Cellular component **C.** Molecular Functions. **D.** Top 20 enriched KEGG pathways. For A-C, the green arcs on the outside parallel to the main chord diagram denote possible (hierarchical) clusters. The number on an arc-node represents the percentage of common genes between two nodes/clusters. The grey links inside the chord diagram connect pairs of GO term/clusters, and indicates the existence of common genes between them.

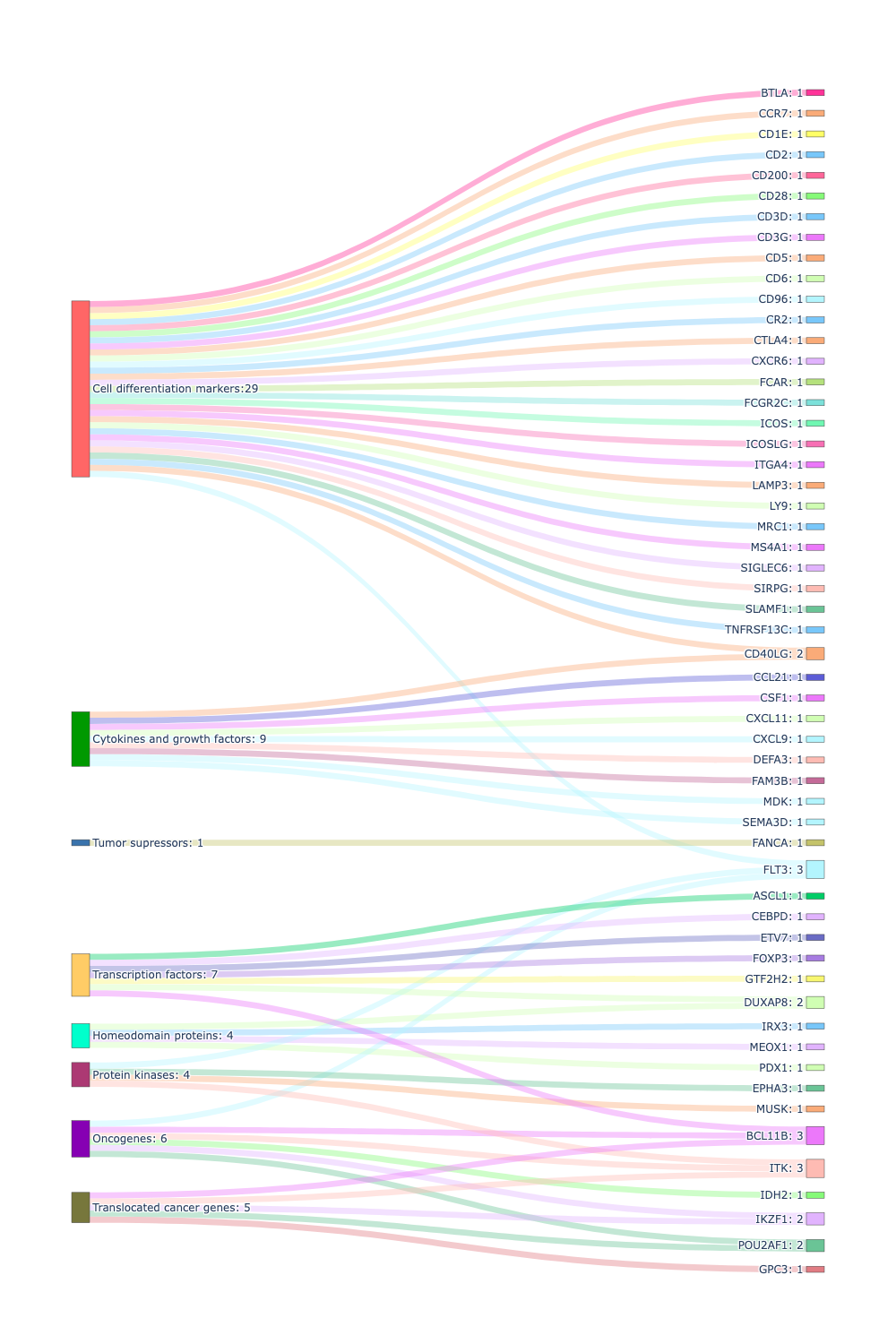

**Figure S7**. Distribution of upregulated genes from HCV+ vs Control group into different gene families in Sankey plot. The number written next to each gene family indicates the number of genes belonging to that gene family. The number next to gene names denotes the number of gene families to which that gene belongs.

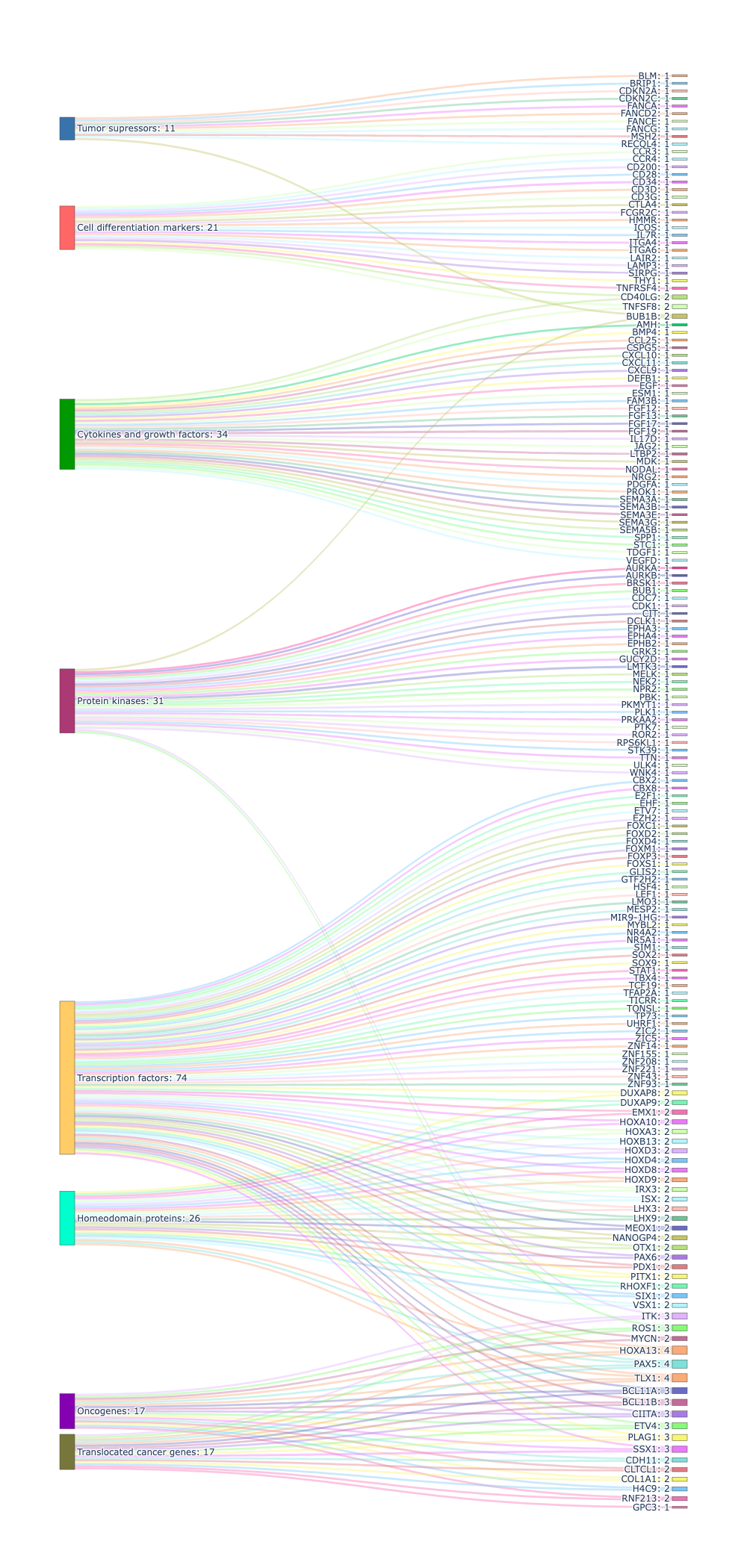

**Figure S8**. Distribution of upregulated genes from HCV+HCC+ vs Control group into different gene families in Sankey plot. The number written next to each gene family indicates the number of genes belonging to that gene family. The number next to gene names denotes the number of gene families to which that gene belongs.

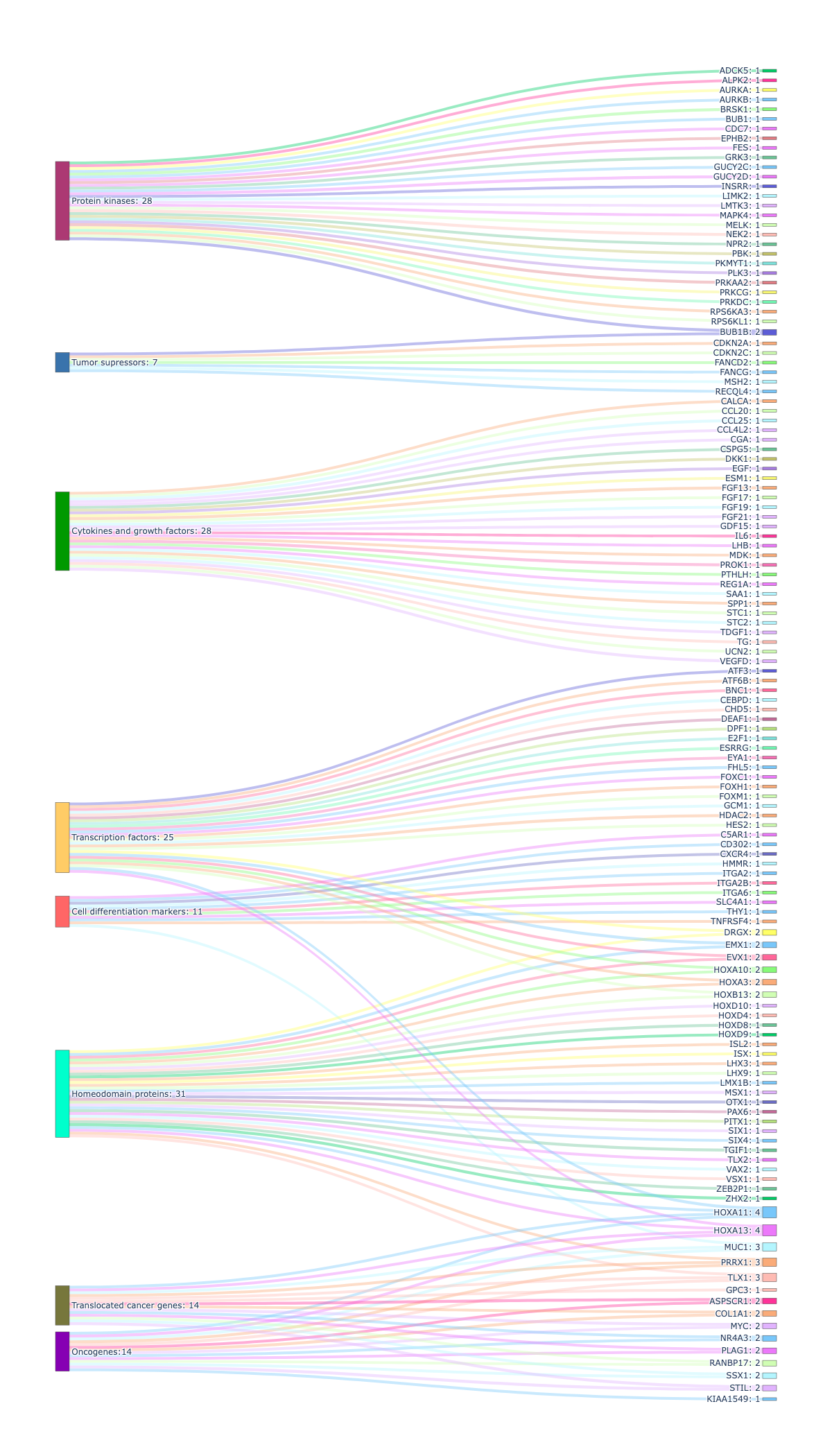

**Figure S9**. Distribution of upregulated genes from HCV+HCC+ vs HCV+ group into different gene families in Sankey plot. The number written next to each gene family indicates the number of genes belonging to that gene family. The number next to gene names denotes the number of gene families to which that gene belongs.

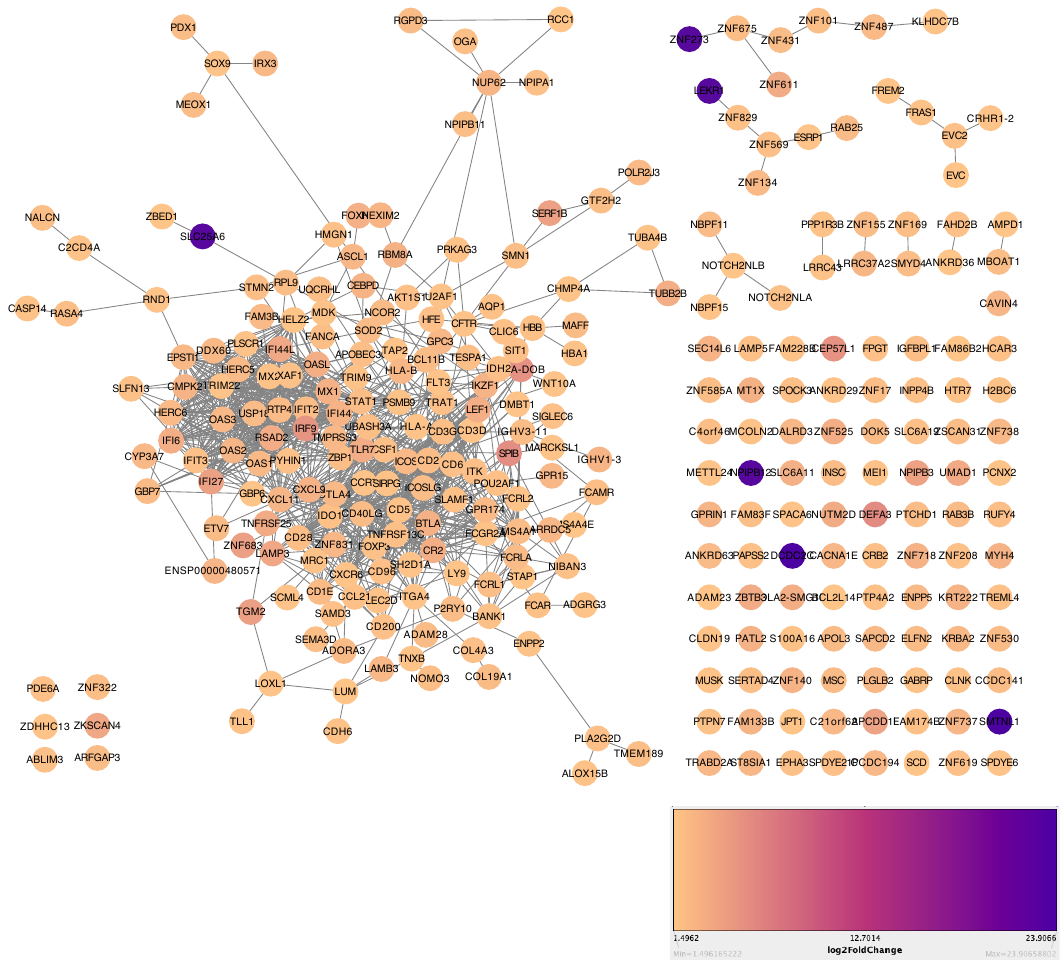

**Figure S10**. Protein-protein interaction network of upregulated genes in HCV+ vs Control. The node colors are shown on the basis of log2 fold change provided in legend.

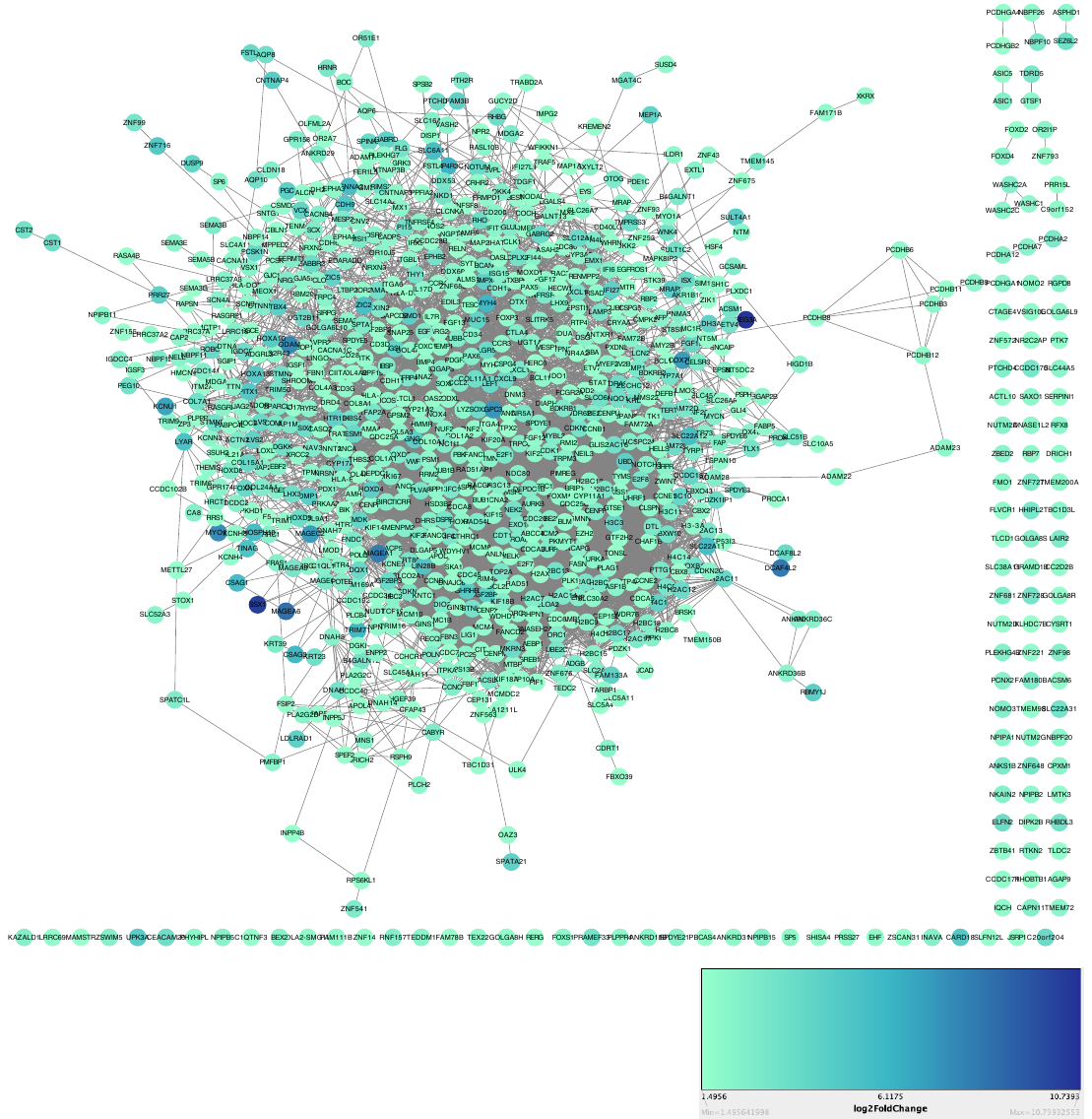

**Figure S11**. Protein-protein interaction network of upregulated genes in HCV+HCC+ vs Control. The node colors are shown on the basis of log2 fold change provided in legend.

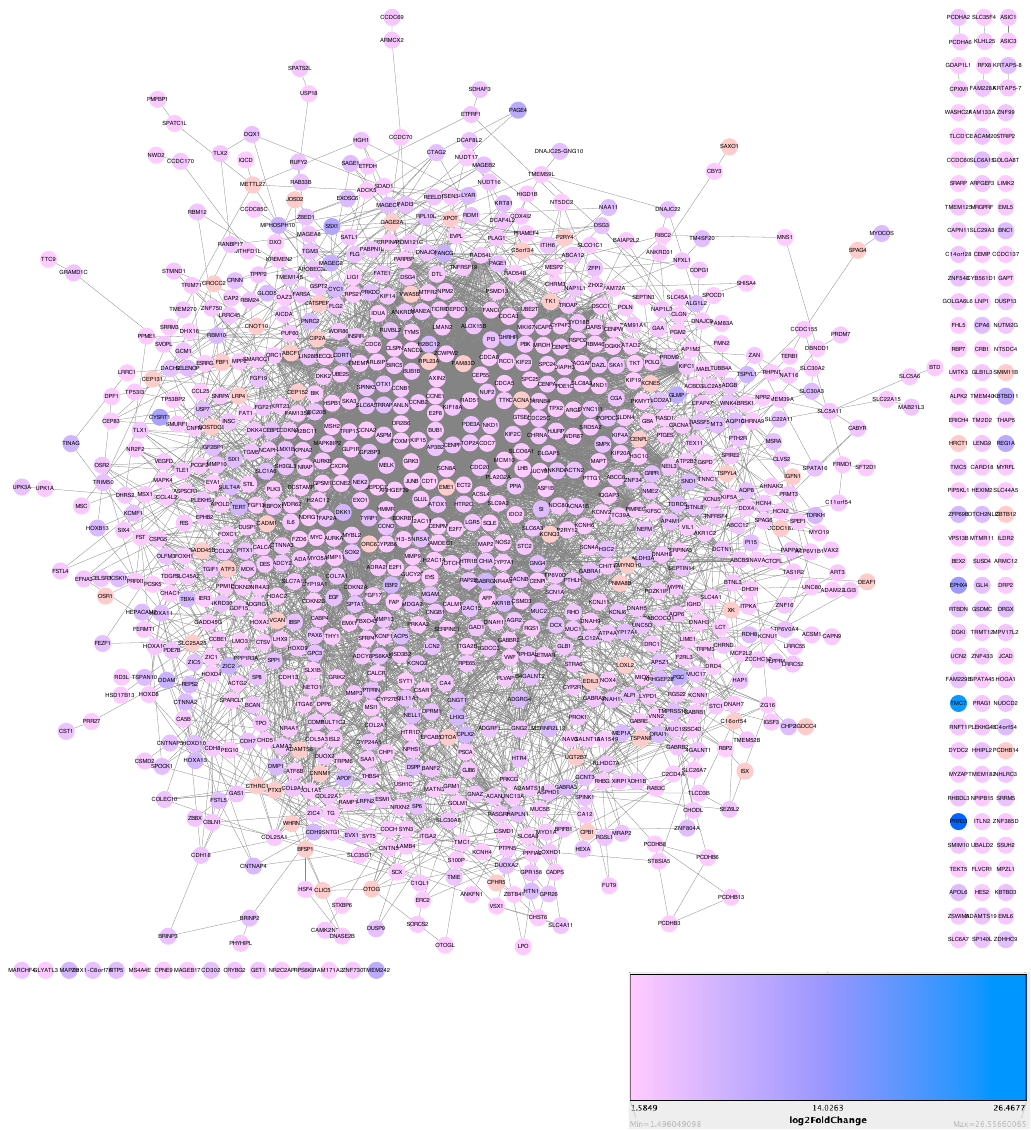

**Figure S12**. Protein-protein interaction network of upregulated genes in HCV+HCC+ vs HCV+. The node colors are shown on the basis of log2 fold change provided in legend.

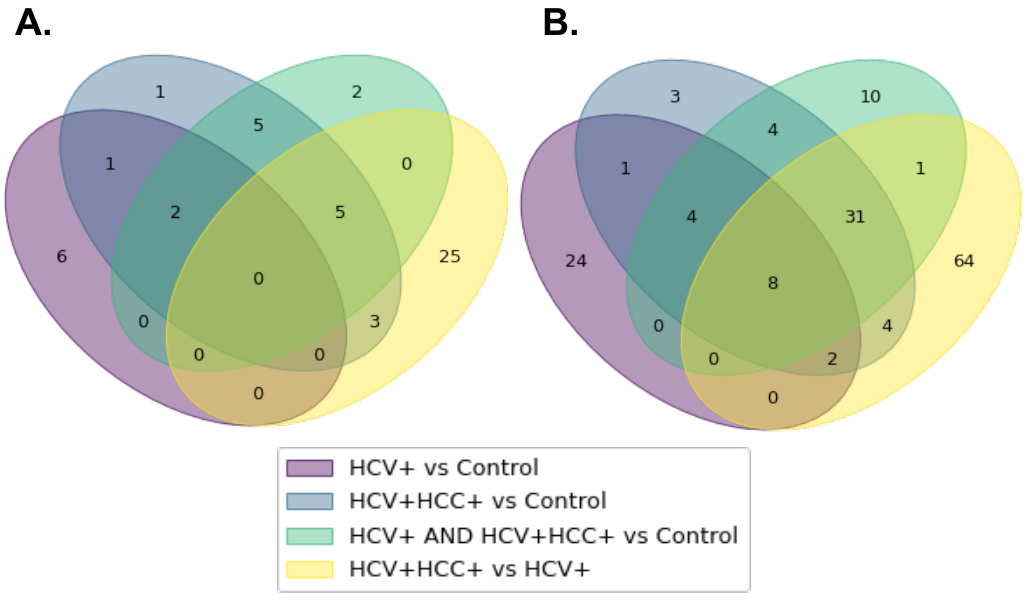

**Figure S13** Venn diagram showing **A.** Overlapping target genes in four studied groups **B.** Overlapping prioritized repurposing drugs among four studied groups. Colors depicting the four studied groups: HCV+ vs Control, HCV+HCC+ vs Control, HCV+ AND HCV+HCC+ vs Control, and HCV+HCC+ vs HCV+ are shown in legend.

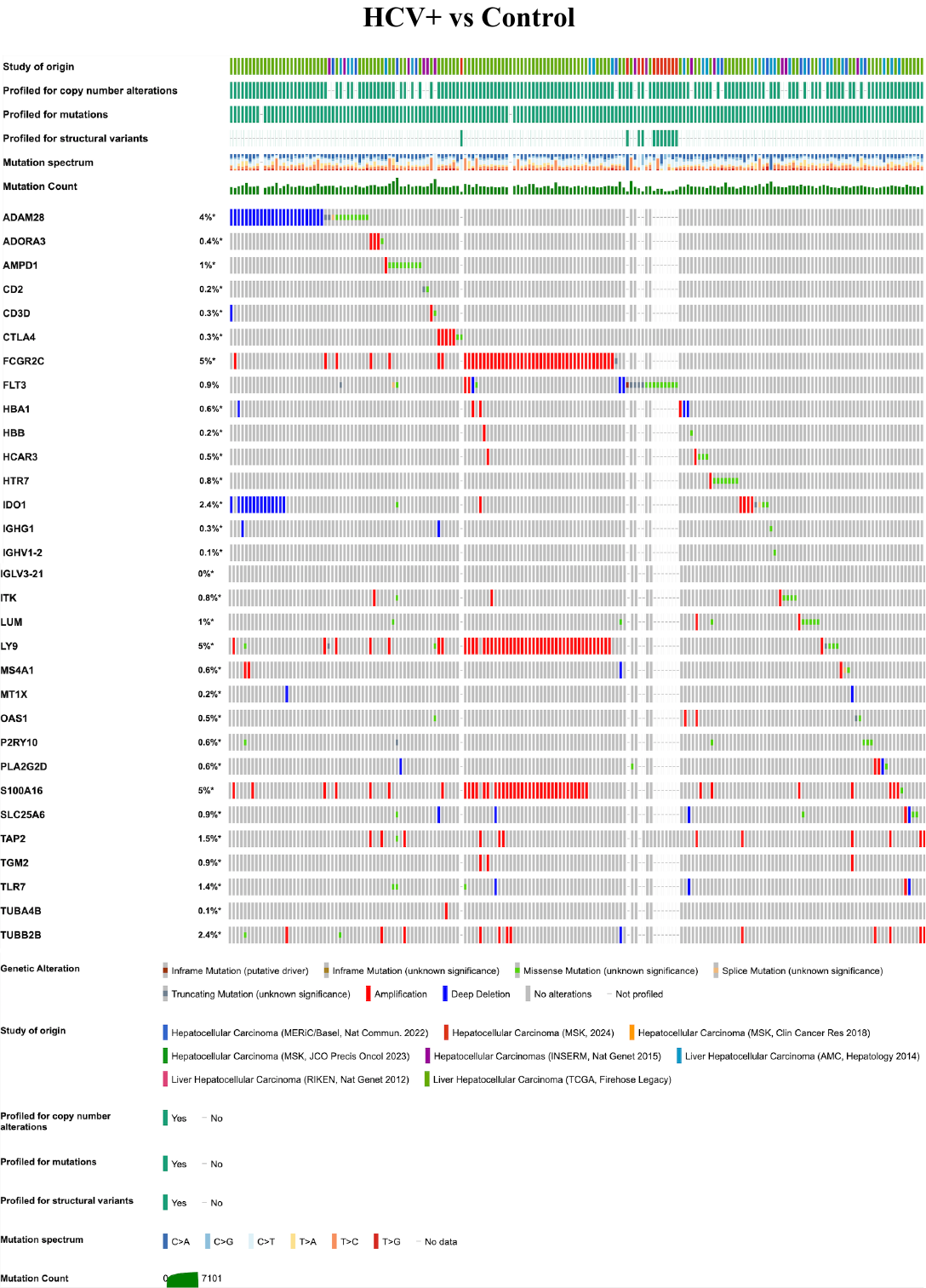

**Figure S14.** Oncoprint image showing the mutational profiles of 31 target genes from HCV+ vs Control group

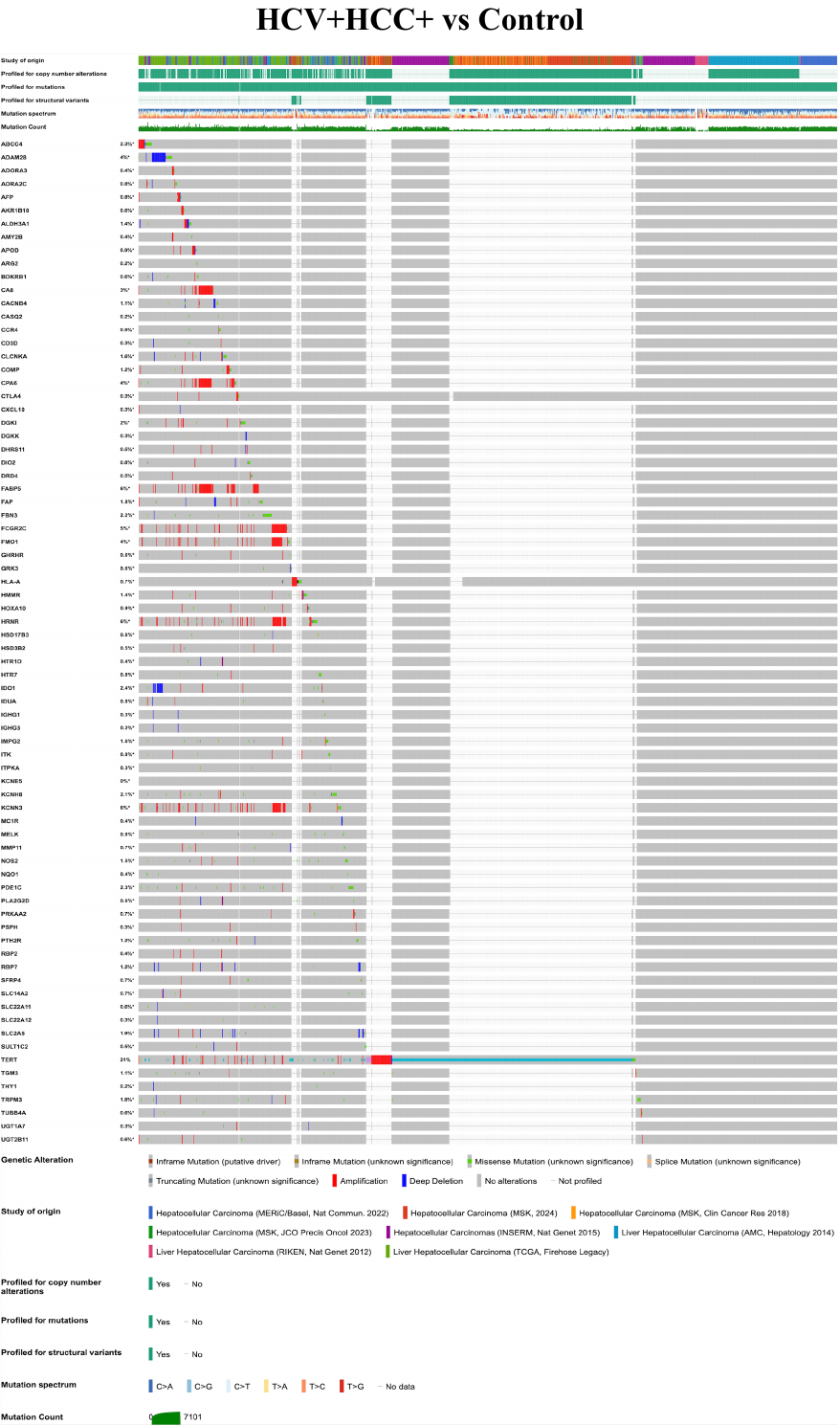

**Figure S15.** Oncoprint image showing the mutational profiles of 76 target genes from HCV+HCC+ vs Control group

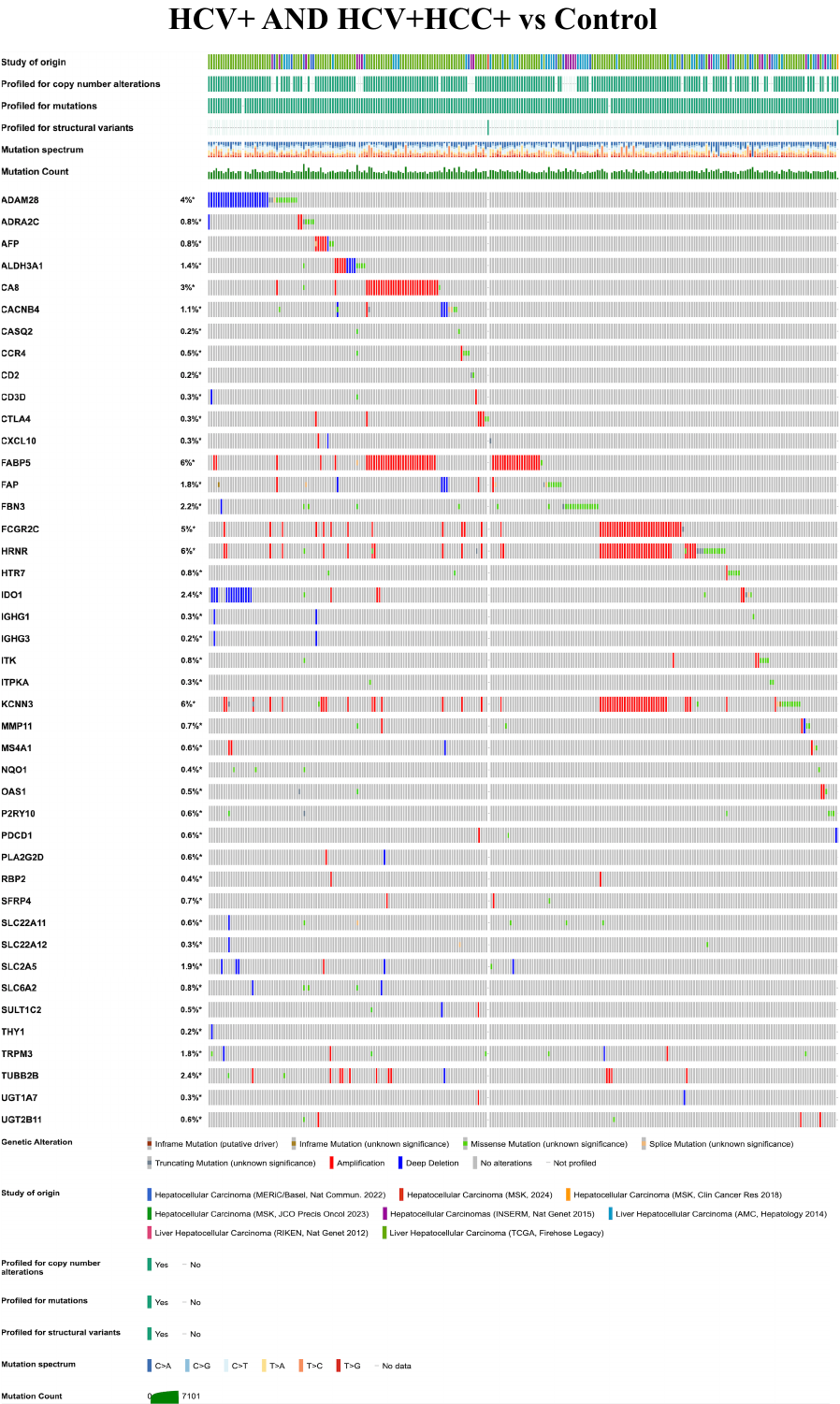

**Figure S16.** Oncoprint image showing the mutational profiles of 43 target genes from HCV+ AND HCV+HCC+ vs Control group

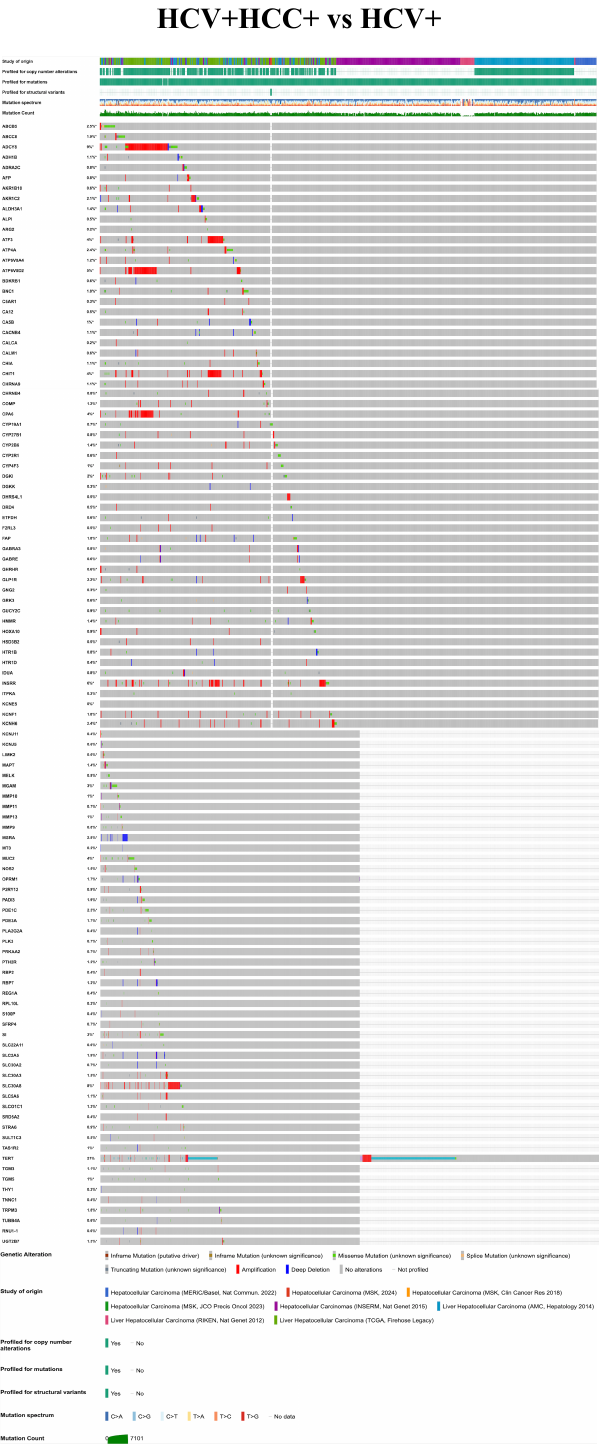

**Figure S17.** Oncoprint image showing the mutational profiles of 109 target genes from HCV+HCC+ vs HCV+ group
